# ‘Cell painting’ with amphiphile-protein conjugate vaccines drives B cell activation and germinal center priming to enhance humoral immunity

**DOI:** 10.64898/2026.09.10.750419

**Authors:** Daman Yadav, Erin L. Templeton, Kathryn Jans, Madison L. Seefeld, Brandon Hu, Justin M. Lehtinen, Noah Sinclair, Brittany L. Hartwell

## Abstract

Most licensed vaccines against infectious diseases mediate protection by priming B cells and humoral immune responses, where efficacy largely relies on activation of germinal centers (GCs) in lymphoid tissues that serve as the training ground for a high affinity antibody response. While subunit vaccines offer safety advantages, they tend to be poorly immunogenic with weak and short-lived antibody responses, presenting a need for engineering strategies that enhance immunogenicity. Amphiphile-protein vaccines, consisting of protein antigens modified with an albumin-binding lipid tail via polyethylene glycol (PEG) linker, hitchhike on albumin following subcutaneous injection to enhance lymphatic trafficking and antigen-specific immune activation. ‘Amph-vaccines’ also demonstrate an ability to insert their lipid tail into cell membranes, effectively ‘painting’ cells with multivalent antigen. We hypothesized that ‘cell painting’ may play a role in driving humoral immunity by sustaining antigen persistence in lymph nodes and generating multivalent antigen presentation for B cell activation. Here, we investigated how amph-vaccine molecular properties (antigen MW and PEG linker length) influenced conformational behaviors (micelle formation, albumin hitchhiking, and membrane insertion) along with the resulting humoral immune response. Using HIV env proteins as model antigens (eOD monomer and MD39 trimer), we investigated cell membrane insertion and B cell activation via calcium flux assay *in vitro*, followed by lymphatic trafficking and immunogenicity studies *in vivo*. Our results demonstrate that amphiphile conjugation is an effective strategy for enhancing humoral immunogenicity of both monomer and trimer protein antigens, where cell painting plays a significant role in driving B cell, GC, and humoral immune activation.

## INTRODUCTION

### Vaccine Kinetics

Most licensed vaccines against infectious diseases mediate protection through B cell activation and priming of humoral immune responses. The magnitude, specificity, and affinity of a humoral response is determined by the germinal center (GC) reaction in lymphoid tissues, such as peripheral lymph nodes (LNs) and mucosal-associated lymphoid tissues (MALT), where antigen-specific immune responses are orchestrated (*1*). Activation of robust GC responses is associated with more effective cross-neutralization of viral variants against pathogens such as SARS-CoV-2 (*2–6*), and is an essential precursor for activating immunity against difficult pathogens like HIV. Upon antigen exposure, B cells within GCs are trained for a high affinity antibody response, undergoing iterative rounds of antigen presentation, somatic hypermutation, and selection to generate high-affinity antibodies, ultimately differentiating into long-lived plasma cells or memory B cells (*7–9*). In this way, GCs serve as a training ground for a high-affinity antibody response and immune memory. Development of effective vaccine strategies, therefore, largely relies on the ability of a vaccine to effectively activate B cells and induce a sustained GC response.

To engineer more effective vaccines, vaccine properties can be tuned to enhance the GC response, both by tailoring vaccine kinetics to enhance trafficking to and persistence in lymphoid tissues and by tailoring antigen multivalency to enhance B cell activation. Vaccine kinetics refers to the spatiotemporal pattern of antigen and adjuvant exposure, where both timing and location are important for the resulting immune response (*7, 10, 11*). Studies have shown that GCs are highly sensitive to the timing and duration of antigen exposure. For example, conventional subcutaneous bolus injections typically result in burst release of antigen that is rapidly cleared from the injection site and draining lymph nodes (dLNs), leading to transient GC reactions and suboptimal antibody titers. In contrast, ‘slow-release’ or ‘escalating-dose’ regimens outperform traditional bolus injections by *prolonging* antigen exposure, as persistent antigen levels sustain GC reactions to promote the maturation of broad high affinity antibodies against antigenic epitopes (*7, 8, 10, 12–14*). Furthermore, spatial kinetics, or the localization of antigen within lymphoid tissue, also influences antigen fate and immune processing. By localizing vaccine components to specialized follicular niches in lymphoid organs, antigens are protected from clearance and degradation, allowing for repeated rounds of B cell signaling that lead to somatic hypermutation and maturation of high affinity antibodies (*15*). In addition to kinetics, antigen multivalency is another critical determinant of vaccine potency. Compared to monovalent antigen, multivalent antigen exhibits higher B cell receptor (BCR) binding affinity through the avidity effect, triggering BCR clustering and BCR-mediated intracellular signaling to enhance B cell activation (*1, 16–20*). By increasing BCR-mediating binding and activation, multivalent antigen directly governs the quality of the GC reaction and is a key driver in generating enhanced and sustained antibody responses.

### Molecular Mechanisms: albumin hitchhiking and cell painting

Most subunit vaccines are administered as conventional bolus injections of soluble protein antigens co-delivered with traditional adjuvants like aluminum salts to induce protective antibody responses (*13*). Yet standard formulations can suffer from rapid antigen clearance from the injection site and inefficient drainage into the lymphatic system, limiting the magnitude and durability of the germinal center response. For example, subunit vaccines containing peptide and small soluble protein antigens (in the absence of particulate carriers / adjuvants) typically drain into systemic circulation following subcutaneous or intramuscular injection due to their small particle size and MW, where they are rapidly diffused, proteolytically degraded, and cleared from the body. Faced with poor pharmacokinetics and inefficient entry into the lymphatic system, subunit vaccines thus tend to be limited by weak immunogenicity and short-lived immune responses. However, the major blood protein albumin, which functions *in vivo* as a fatty acid transporter, has been exploited as a drug delivery chaperone protein to target both small molecule drugs and subunit vaccine cargo to LNs following injection. Albumin preferentially traffics into the lymphatics from interstitial space given its MW (∼67 kDa) (*7, 10, 19, 21, 22*) (*23–27*). Albumin is also transcytosed by the neonatal Fc receptor (FcRn) expressed on vascular endothelial and epithelial cells, undergoing recycling that protects it from lysosomal degradation and clearance (*24*). Thus, albumin hitchhiking not only enhances drug accumulation in secondary lymphoid organs, but also promotes longer drug and antigen half-life.

Here, we leveraged this endogenous transport pathway by conjugating protein antigens to an albumin-binding lipid tail, synthesizing amphiphile vaccines (‘amph-vaccines’) that bind endogenous albumin following injection to ‘hitchhike’ through the lymphatic vasculature to dLN(s). We evaluated amphiphile conjugates of monomer and trimer protein antigens to target B cells and the humoral immune response. Our amphiphile vaccine platform consists of a protein immunogen covalently conjugated to a diacyl lipid tail (DSPE) via a polyethylene glycol (PEG) linker. This drug delivery strategy was previously demonstrated to enhance LN trafficking of small peptide antigens and molecular adjuvants (CpG) following subcutaneous injection, leading to significantly enhanced cellular immune responses (*12, 28–31*). We previously showed that DSPE-PEG modification also enhances mucosal uptake and mucosal associated lymphoid tissue (MALT) trafficking of monomeric proteins following intranasal administration, leading to significantly enhanced systemic and mucosal humoral immune responses (*1, 21, 22*).

In addition to hitchhiking on albumin, amph-vaccines demonstrate a secondary behavior whereby the lipid tail inserts nonspecifically into cell membranes (*1, 28, 32*). We hypothesized that membrane insertion may impact vaccine efficacy in two ways: 1) by promoting retention and persistence of vaccine antigen in secondary lymphoid tissues, leading to altered vaccine kinetics; and 2) by effectively ‘painting’ surrounding cells with multiple copies of antigen, leading to multivalent antigen presentation and enhanced B cell activation. Rather than relying on uptake and presentation first by other APCs or low avidity monovalent BCR binding to target B cells, we posited that membrane insertion could provide a unique mechanism of multivalent antigen presentation through high avidity engagement of BCRs, leading to BCR crosslinking to enhance activation. Membrane insertion has previously been studied with amphiphile conjugates of small peptides, haptens, and oligonucleotides, such as FITC and CpG – for example, using amph-FITC to decorate the surface of tumor-infiltrating lymphocytes to activate FITC-specific CAR-T cells (*28*), or using membrane anchoring of amph-CpG and amph-peptides within LNs to polarize T cell responses (*12, 32*). However, to our knowledge the contribution of membrane insertion of *amph-protein vaccines on B cell priming and humoral immunity* has remained unexplored and unknown. It was also unclear whether B cell activation from amph-protein vaccines was driven by trafficking alone, or also membrane-mediated multivalent presentation or self-assembly of amph-protein conjugates into micelles in secondary lymphoid tissues. Thus, here we sought to compare multiple versions of amph-protein conjugates with differing conformational properties to their soluble protein controls to investigate the role of platform design and membrane insertion, or ‘cell painting’, on B cell activation and immunity.

### Model System

There is a <u>critical need</u> for development of immunization strategies that induce high affinity and broad humoral immune responses across infectious disease settings. Here, we were interested in testing ‘cell painting’ as a platform strategy for enhancing vaccine efficacy and humoral immune activation using HIV as a model system. HIV poses a particular challenge for vaccination efforts due to its antigenic diversity and immune evasion mechanisms, requiring vaccine strategies that drive broad and durable GC responses capable of maturing broadly neutralizing antibodies (bnAbs) (*1, 33, 34*). For model antigens to test in the amph-vaccine platform, we selected gp120 eOD (engineered outer domain) monomer and MD39 SOSIP trimer. eOD is an engineered version of the gp120 glycoprotein displayed on the HIV viral envelope, designed by the Schief group to have higher binding affinity for the diverse CD4 binding sites that are targeted by VRC01-class bnAbs (*35, 36*). For example, eOD-GT8 60mer (eOD-60mer) is a protein nanoparticle (NP) presenting 60 copies of eOD that is currently being investigated in clinical trials as a priming immunogen to target germline precursor B cells as a strategy to initiate CD4 binding site directed broadly neutralizing antibodies against HIV (*37, 38*). We have previously used eOD as a testbed antigen in DSPE-PEG2K amphiphiles to test intranasal amph-vaccine immunization in mice and nonhuman primates (*1*). However, achieving a functionally protective antibody response against HIV is thought to require subsequent boosts with native-like trimers capable of inducing tier 2 broadly neutralizing antibodies (bnAbs) against conformational epitopes. Trimers such as BG505 SOSIP and its stabilized derivative MD39 were engineered to mimic the native HIV Env spike (*39*). These stabilized trimers have demonstrated the ability to elicit high neutralizing antibody titers in animal models and have successfully protected rhesus monkeys from SHIV challenges (*40–42*). More importantly, these native-like trimer-based immunization strategies have also transitioned into human clinical trials, where they have been shown to successfully induce tier 2 neutralizing antibodies in phase I clinical trials (*43*). Therefore, we selected eOD and MD39 to test within the amph-vaccine platform for cell painting as model antigens that share common epitopes and specificity for VRC01 but differ in their physicochemical properties (MW, stability, conformation) and potential to elicit bnAbs.

Previous work by Zhang *et al* demonstrated that the PEG linker in DSPE-PEG amphiphile conjugates plays a critical role in stabilizing membrane insertion, dependent on PEG linker length; amphiphile-hapten conjugates with PEG linkers of 2000 Da MW exhibited stable cell membrane insertion while PEG MWs of 5000 Da or greater significantly decreased insertion (*28, 32*). We applied this concept here to make a version of amph-eOD with PEG5K to reduce cell membrane insertion. By altering both PEG linker length and antigen MW (eOD monomer vs MD39 trimer) conjugated to DSPE, we synthesized amph-protein constructs with varying levels of membrane insertion while controlling for albumin hitchhiking and micelle formation in order to separately investigate ‘cell painting’ from these other behaviors. First, molecular properties and conformation behaviors were characterized: amph-vaccine micelle formation using DLS particle sizing and TEM, albumin binding using a combination of SEC and affinity chromatography, and cell membrane insertion using *in vitro* flow cytometry and imaging assays. B cell activation was evaluated *in vitro* using a calcium flux assay in an engineered antigen-specific glVRC01 B cell line. Trafficking and persistence in dLNs was measured *in vivo* by IVIS. Lastly, B cell activation was evaluated *in vivo* by measuring GC expansion in dLNs following subcutaneous immunization in mice, with longitudinal and long-lasting humoral immune activation evaluated by ELISA and ELISPOT, respectively. Overall, our goal was to investigate how amphiphile-protein vaccine design parameters (such as antigen MW and PEG linker length) and molecular behaviors (like cell painting or micelle formation) drive B cell and humoral immune activation.

## RESULTS

### Protein amphiphile conjugates exhibit variable micelle formation dependent on antigen molecular weight

To evaluate how amphiphile behaviors like micelle formation and membrane insertion impact B cell activation and vaccine immunogenicity, we first synthesized a set of amphiphile-protein conjugates with different physicochemical and conformational properties that would drive these behaviors (**Fig. 1**). Using HIV envelope (Env) proteins as model antigens, we synthesized amphiphile-protein conjugates with two different antigen molecular weights and PEG linker lengths. First, for a model monomer antigen, we selected HIV Env eOD-GT8: a germline-targeting ∼22 kDa engineered outer domain of gp120 previously shown to prime VRC01-class broadly neutralizing antibody precursors in a phase 1 clinical trial (*35, 36, 38*). The universal T helper epitope ‘PADRE’ (pan human leukocyte antigen DR-binding epitope) was included at the eOD C-terminus. eOD-PADRE (hereafter referred to as eOD) was modified with an N-terminal cysteine, then site-specifically conjugated to maleimide-functionalized PEG2K–DSPE (polyethylene glycol-1,2-distearoyl-sn-glycero-3-phosphoethanolamine) via stable thioether linkage to form ‘amph-PEG2K-eOD’ amphiphile conjugates (**Fig. 1A, S1A,C**). A second eOD amphiphile conjugate was synthesized using a longer PEG5K spacer to form ‘amph-PEG5K-eOD’ as a control for cell painting (**Fig. 1B**), motivated by prior work that showed amphiphile conjugates with longer PEG linkers (≥5K) exhibited markedly reduced membrane insertion (*32*). *Amph-PEG5K-eOD was thus included in these studies as an amphiphile control to determine the role of cell painting, as a conjugate with equivalent micelle formation and albumin binding behavior as amph-PEG2K-eOD but <u>significantly reduced cell painting</u>*. Indeed, both amph-PEG2K-eOD and amph-PEG5K-eOD formed stable ∼25 nm micelles in the absence of albumin compared to ∼5 nm unmodified eOD protein, as shown by SEC (**Fig. 1C**), DLS (**Fig. 1D-E**), and TEM (**Fig. 1I**). Micelle formation of amph-PEG2K-eOD and amph-PEG5K-eOD was concentration dependent, with stable micelles forming above a critical concentration (CMC) of around 100 nM (**Fig. 1E**).

**Figure 1.**
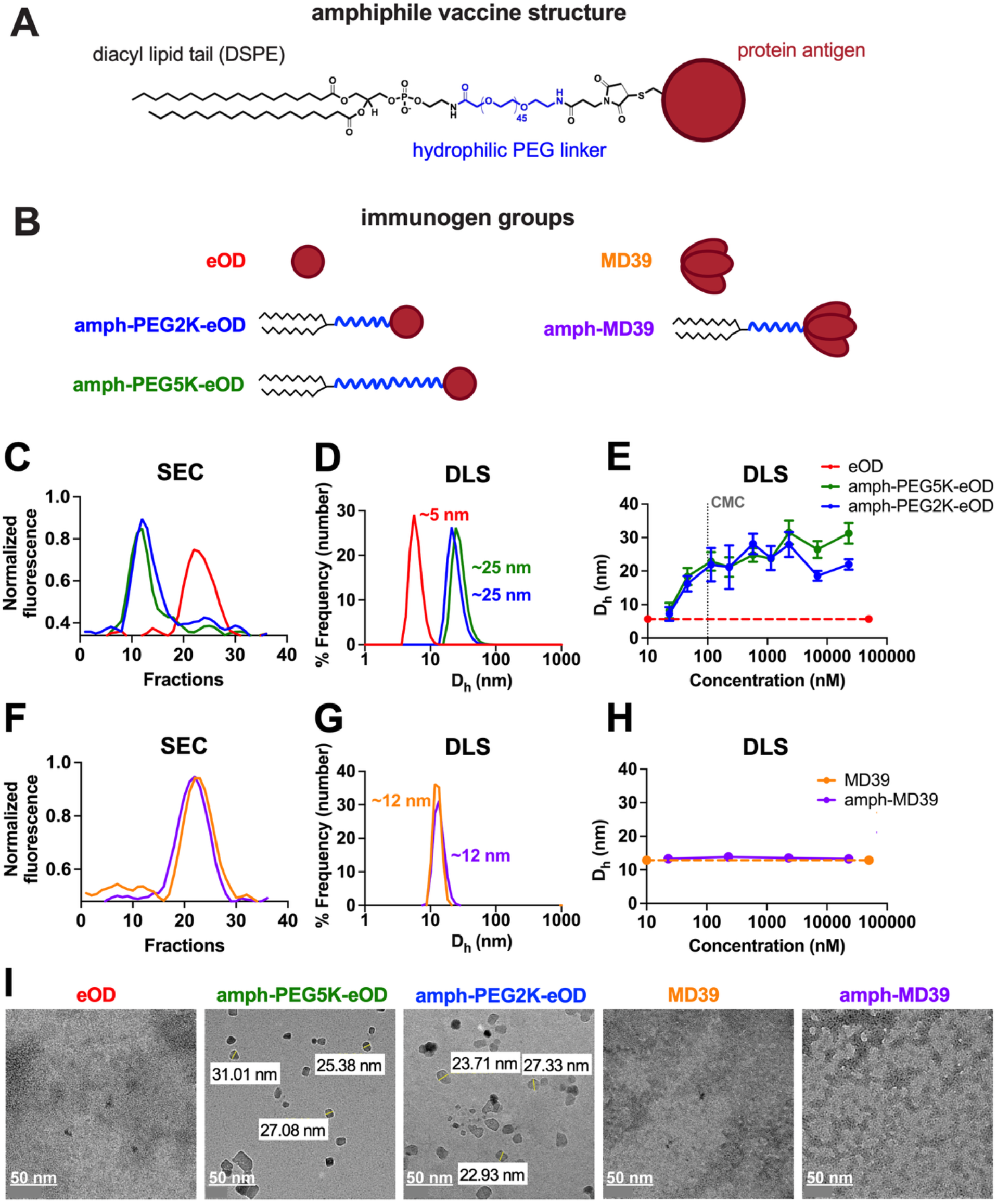
Synthesis and physicochemical characterization of amphiphile-protein conjugates. **A)** Amphiphile vaccine structure, consisting of protein antigen cargo conjugated to a lipophilic diacyl tail (DSPE) via a hydrophilic polyethylene glycol (PEG) linker. **B)** Representative structures of eOD and MD39 immunogen groups, including soluble controls and amphiphile conjugates. **C-E)** Characterization of eOD versus amph-PEG5K-eOD and amph-PEG2K-eOD: **C)** Size exclusion chromatography (SEC) elution profiles; **D)** Dynamic light scattering (DLS) analysis, showing hydrodynamic diameter (D_h_) as number-weighted % frequency; **E)** DLS particle size as a function of concentration to determine critical micelle concentration (CMC) for eOD amph-conjugates. **F-H)** Characterization of MD39 versus amph-PEG2K-MD39: **F)** SEC elution profiles; **G)** DLS analysis, shown as number-weighted % frequency; **H)** DLS particle size as a function of concentration for MD39 amph-conjugate. **I)** Representative transmission electron microscopy (TEM) images of all immunogen constructs.

Second, for a model trimer antigen we selected HIV Env BG505 MD39 SOSIP: a conformationally dependent native-like trimer of ∼220 kDa (depending on glycosylation state) that is a lead immunogen candidate for eliciting broadly neutralizing antibodies (bnAbs) against HIV in nonhuman primates and humans (**Fig. 1B, S1B**). As a larger and more structurally complex protein than eOD monomer, MD39 allowed us to test the role of antigen size on the amphiphile platform’s functionality. MD39 was modified with a C-terminal cysteine, then first conjugated to an intermediate PEG4-dibenzocyclooctyne linker (maleimide-DBCO-PEG4) before conjugating with PEG2K-DSPE-azide through copper-free click chemistry **(Fig. S1D)** (*44, 45*). Using an intermediate DBCO linker to account for steric hindrance of the larger MD39 trimer enabled efficient amphiphile modification of the trimer protein (**Fig. 1B**). In contrast to amph-eOD conjugates, amph-MD39 was sterically hindered from forming micelles in the absence of albumin, maintaining a particle size of ∼12 nm that was equivalent to that of unmodified MD39 as demonstrated by SEC (**Fig. 1F**), DLS (**Fig. 1G-H**), and TEM (**Fig. 1I**). Amph-MD39 maintained this particle size independent of concentration (**Fig. 1H**). We hypothesized that amph-MD39 would exhibit albumin binding to a similar extent as amph-eOD, thus inclusion of this group allowed us to test the role of micelle formation for B cell activation.

### Protein amphiphile conjugates exhibit albumin binding independent of PEG linker length or protein MW

Previously we showed that DSPE-PEG2K conjugates of protein monomers exhibited albumin binding and ‘hitchhiking’ to alter trafficking in vivo (*1*). DSPE-PEG2K conjugates bind to albumin with an equilibrium dissociation constant (K_D_) of ∼125 nM (*^12^*). To evaluate whether albumin binding differed with DSPE-PEG conjugates incorporating a longer linker (PEG5K compared to PEG2K) or larger protein (MD39 trimer compared to eOD monomer), we tested the ability of all protein-amphiphiles to bind albumin via lipid tail while confirming that structure and antigenicity of the conjugated proteins were conserved (**Fig. 2**). First, albumin binding was evaluated in vitro by incubating fluorescently labeled protein immunogens with albumin-functionalized agarose resin for 2 hours at 37C, followed by separation of resin and measurement of fluorescent protein that remained bound to resin versus eluted in the filtrate. Proteins (600 nM) were incubated with a molar excess of BSA (100 µM) to ensure protein binding to albumin was not saturated (*46*). To ensure a consistent and accurate signal, resin trapped within the filter membrane was excluded from the analysis. Compared to unmodified eOD and MD39, both amph-eOD and amph-MD39 conjugates exhibited significantly higher binding to albumin, with over 70% of the conjugated antigens retained on the resin compared to only ∼10% of the unmodified soluble proteins (**Fig. 2A, B**). No significant difference was observed between amph-PEG2K-eOD and amph-PEG5K-eOD, or between amph-PEG2K-eOD and amph-MD39, demonstrating similar albumin binding independent of PEG linker length or protein MW. Furthermore, we also evaluated AF647 fluorescence in the filtrate fractions and observed a similar trend, with both soluble proteins (eOD and MD39) significantly showing up in the eluant, indicating no binding to direct binding to albumin **(Fig. S2)**.

**Figure 2.**
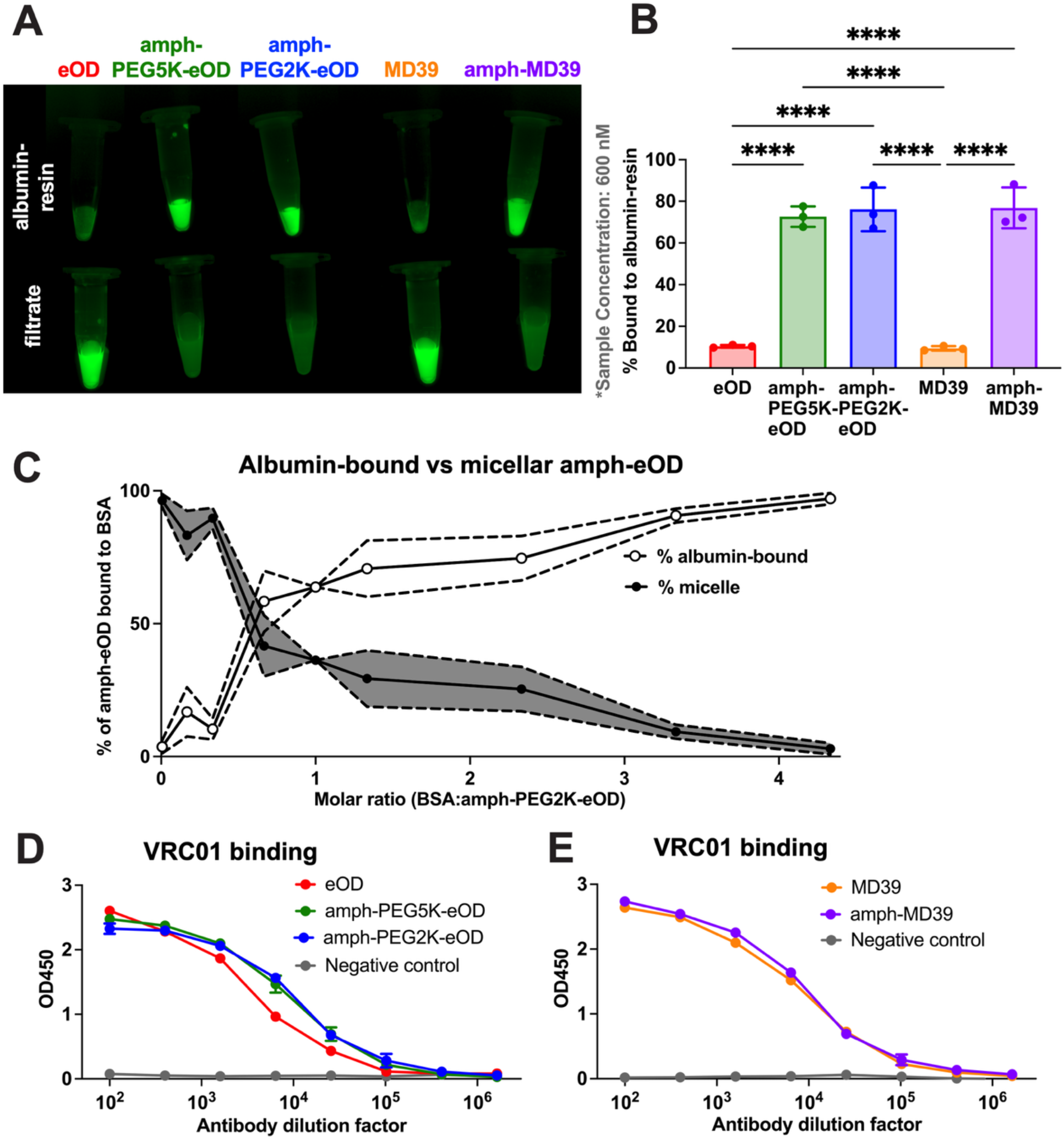
Amphiphile conjugation of eOD and MD39 protein antigens enables albumin binding while conserving antigenicity. **A)** AF647 fluorescence image on gel imager shows retention of AF647-protein vs AF647-amph-protein on albumin-conjugated agarose resin by affinity chromatography following a 1 hr incubation at 37 °C. **B)** Percent of protein/amph-protein bound to albumin-resin following 1 hr incubation at 37 °C, quantified from fluorescence signal in (A). **C)** Percentage of amph-PEG2K-eOD conjugate in micellar form versus albumin-bound state following incubation with bovine serum albumin (BSA) at different molar ratios for 1 hr at 37 °C, determined by measuring AF647-eOD and AF488-BSA fluorescence across all collected SEC fractions. **D)** Relative antigenicity of eOD and **E)** MD39 constructs based on binding with hVRC01 antibody compared to PBS negative control. Statistical significance determined by one-way ANOVA followed by Sidak’s post-hoc test. Data showing mean ± SEM (n=3).

To further characterize albumin binding and to quantify the relative amount of amphiphile that was albumin-bound versus micellar as a function of BSA concentration, a co-elution assay was performed using SEC. Excess AF647-labeled amph-PEG2K-eOD (1500 nM, selected above the CMC to ensure stable micelle formation in the absence of albumin) was incubated with increasing concentrations of AF488-labeled BSA ranging from 250 nM to 6500 nM, then separated by SEC to determine whether amph-eOD protein eluted with micelle or albumin fractions (**Fig. 2C**). The BSA range was selected to transition from a sub-stoichiometric molar ratio of BSA:amphiphile to a concentration ratio that more closely mimicked that found *in vivo* in the interstitial space. As hypothesized, we observed an inverse relationship between amph-eOD albumin binding and micelle formation. As albumin concentration increased, the fraction of amph-eOD bound to albumin also increased while the micellar fraction decreased, resulting in a concentration-dependent shift toward albumin-bound fractions. At a 1:1 molar ratio of BSA:amph-eOD, around 70% of amph-eOD was bound to albumin. *In vivo*, this stoichiometry is expected to shift further to the right as amphiphile is diluted upon dissemination into the interstitial space and lymphatics while albumin concentrations reach greater excess. Based on conservative mean estimates of albumin concentrations in interstitial space (∼10 g/L or ∼150 µM)(*47–49*) compared to our injected concentration of amph-eOD (5 µg / 100 µl or 2.3 µM), we estimate greater than 60-fold molar excess of albumin compared to amphiphile in the interstitial space (*29*). Thus, our findings indicate that amph-vaccines exhibit concentration-dependent binding to albumin and are most likely to exist in the non-micellar form *in vivo*.

Lastly, the antigenicity of all conjugates (binding of antigen to Env-specific VRC01 antibody) was assessed by ELISA. No significant differences in antigenic epitope binding were observed between eOD, amph-PEG2K-eOD, and amph-PEG5K-eOD (**Fig. 2D**), or between MD39 and amph-MD39 (**Fig. 2E**), indicating that amphiphile modification did not alter key structural epitopes of either protein.

### Protein amphiphile conjugates exhibit variable ‘cell painting’ (membrane insertion) dependent on PEG linker length

In addition to their albumin hitchhiking behavior, amphiphile vaccines demonstrate a secondary mechanism of **‘cell painting’** whereby the amphiphile tail nonspecifically inserts into cell membranes to effectively ‘paint’ cells with antigen (**Fig. 3**). Membrane insertion behavior has been demonstrated previously by us and others with amphiphile-peptide or -hapten (*28, 32*), amphiphile-adjuvant (*12, 50*), and amphiphile-protein conjugates (*1*). This prior work had shown that membrane insertion with amphiphile-peptide/hapten conjugates was dependent on PEG linker length (*32*) with longer PEG linkers (≥5K) destabilizing membrane anchoring, but this dependence had not yet been investigated with amphiphile conjugates of larger proteins. So, we next wanted to determine if cell painting with amph-protein conjugates was dependent on PEG linker length or protein size (i.e., inhibited by incorporation of larger trimers due to steric hindrance).

**Figure 3.**
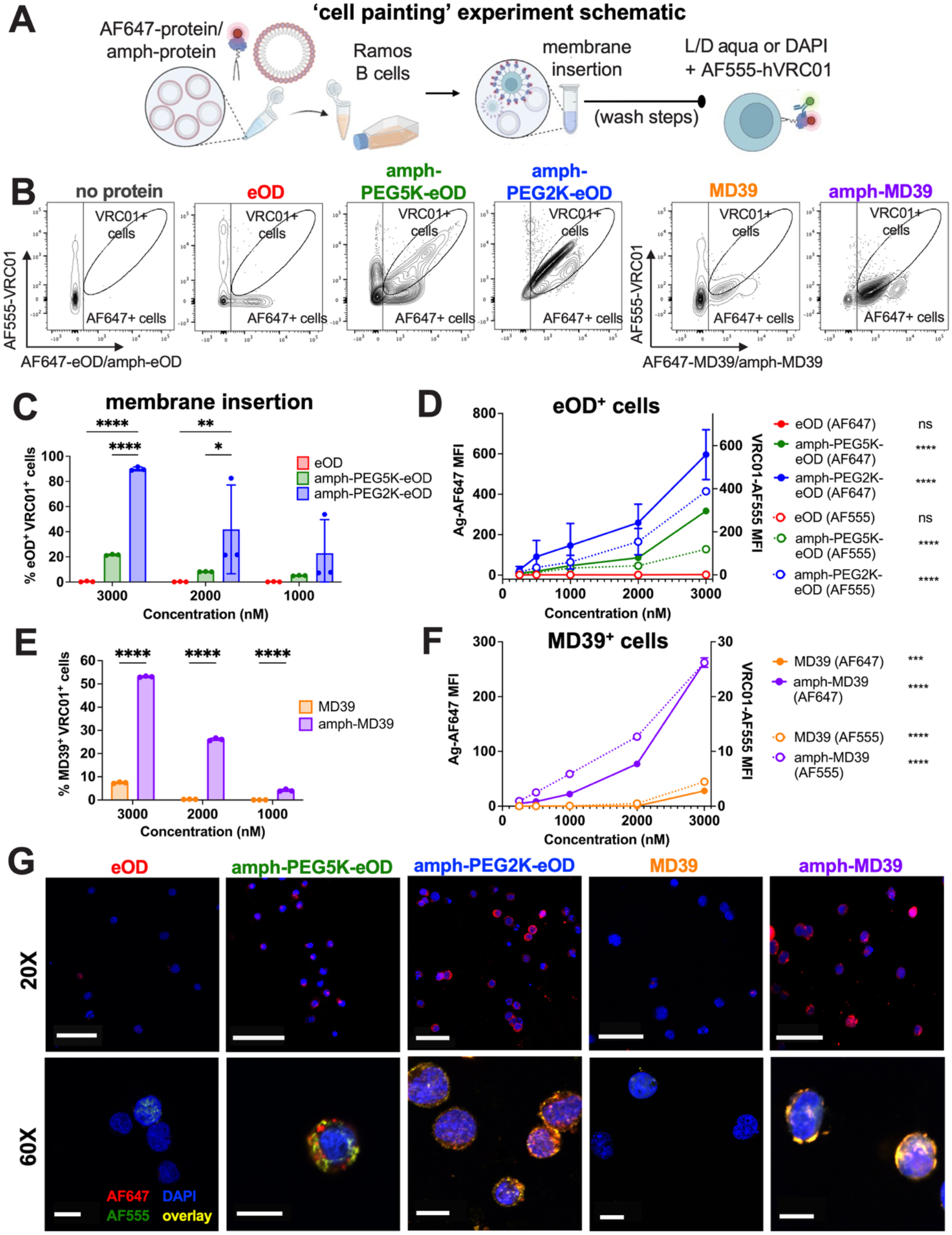
Amphiphile-PEG2K conjugates of eOD and MD39 exhibit lipid tail membrane insertion to facilitate ‘cell painting’ with antigen. A) Experiment schematic for membrane insertion assay: Ramos B cells were incubated with AF647-labeled protein or amph-protein conjugates in cRPMI for 1 hr at 37°C, washed to remove unassociated protein, then stained with L/D aqua (for flow cytometry) or DAPI (for confocal microscopy) and AF555-hVRC01 Env-specific antibody to detect eOD or MD39 Env proteins on the cell surface. **B)** Representative flow cytometry plots of AF647-eOD or -MD39 immunogen and VRC01 binding in Ramos B cells when incubated at 3000 nM. AF647^+^VRC01^+^ double-positive cell gate represents ‘cell painting’. **C-D)** Cell painting with eOD immunogens: **C)** Percentage of cells double-positive for eOD and VRC01 quantified by flow cytometry; statistical significance determined by two-way ANOVA followed by Tukey’s post-hoc test. **D)** Mean fluorescence intensity (MFI) of eOD/amph-eOD and VRC01 as a function of eOD concentration; statistically significant nonzero slope determined by simple linear regression. **E-F)** Cell painting with MD39 immunogens: **E)** Percentage of cells double-positive for MD39 and VRC01 quantified by flow cytometry; statistical significance determined by unpaired t-test. **F)** MFI of MD39/amph-MD39 and VRC01 as a function of MD39 concentration; statistically significant nonzero slope determined by simple linear regression. All data shown are presented as mean +/-SEM (n=3). **G)** Representative images obtained by confocal microscopy show localization of AF647-labeled immunogens (red) in Ramos B cells stained with DAPI (blue) and colocalization with AF555-VRC01 (green) on the cell membrane in amph-PEG2K-eOD and amph-MD39 incubated cells. Scale bars equal 50 µm (20X) and 10 µm (60X).

To evaluate ‘cell painting’ behavior with different amphiphile-protein conjugates, we measured cell membrane insertion by flow cytometry and fluorescence microscopy (**Fig. 3**). First, AF647-labeled soluble proteins or amph-protein conjugates were incubated with nonspecific Ramos B cells in serum-containing media for 1 hour at 37C, then washed to remove any excess protein unassociated with the cell membrane. Next, cells were stained with AF555-labeled hVRC01, an Env-specific antibody that binds eOD and MD39 proteins, to distinguish between antigen decorating or ‘painting’ the cell surface from antigen that was internalized (**Fig. 3A**). Membrane insertion was quantified using flow cytometry by incubating different concentrations of protein conjugates, and measuring AF647-protein/amph-protein and AF555-hVRC01 fluorescence, with membrane insertion represented by AF647^+^AF555^+^ double positive cells (**Fig. 3B, S3A,B)**. Double positive cell population is representative of amphiphile painted cells that have the conjugate being anchored in the cell membrane expressing AF647-protein signal and the bound AF555-hVRC01 antibody signal.

DSPE-PEG2K amphiphile conjugates of eOD and MD39 exhibited significantly greater cell-painting relative to unmodified proteins (**Fig. 3B-F**). Amph-PEG2K-eOD showed significantly greater cell painting (cells double positive for eOD and VRC01) at 3000 nM and 2000 nM compared to unmodified eOD (*p*<0.0001 and *p*<0.01, respectively), with over 90% of cells painted following incubation at 3000 nM – a ∼90-fold increase over soluble eOD, which showed negligible signal on the cell surface (**Fig. 3BC**). Similarly, amph-MD39 showed significantly greater cell painting (cells double positive for MD39 and VRC01) at all concentrations tested compared to unmodified MD39 (*p*<0.0001), with over 50% of cells painted following incubation at 3000 nM, 30-fold and 10-fold enhancement in % cell painting by amph-MD39 conjugates compared to MD39 in 2000 and 1000 nM concentrations respectively (**Fig. 3E**). These data demonstrate that larger proteins are still capable of membrane insertion if conjugated to a lipid tail. Furthermore, comparing different amph-eOD conjugates revealed that PEG spacer length significantly influenced membrane association, as cell painting was markedly reduced in amph-eOD conjugates incorporating a longer PEG5K linker . At 3000 nM, ∼90% of cells were painted with amph-PEG2K-eOD compared to just ∼20% with amph-PEG5K-eOD – roughly four-fold reduction in membrane insertion with the PEG5K variant across all concentrations (*p*<0.0001 at 3000nM; *p*<0.05 at 2000nM) (**Fig. 3C, S3B**). In addition to the percentage of double-positive cells, the mean fluorescence intensity (MFI) also increased in a concentration-dependent manner, indicating that cell painting is characterized by increased density of membrane-anchored antigen per cell in addition to increased number of cells labeled. (**Fig. 3D, F**). Lastly, confocal microscopy of fixed, stained cells supported the flow cytometry observations, showing colocalized AF647-protein and AF555-VRC01 in amph-PEG2K-eOD and amph-MD39 incubated cells, but negligible AF647 or AF555 surface signal in eOD or MD39 incubated cells (**Fig. 3G**). In painted cells, fluorescence was predominantly localized to the cell membrane and equally distributed across the surface **(Fig. S3C)**.

### Amphiphile-mediated micelle formation and membrane insertion both enhance B cell activation *in vitro*

We hypothesized that multivalent antigen presentation, either in the form of amph-protein micelles or cells ‘painted’ with membrane anchored amph-protein, would enhance B cell activation compared to soluble / monovalent protein antigen. Thus, we performed calcium flux assays using a gene-recombinant glVRC01 B cell line that expresses B cell receptors (BCRs) specific for HIV antigens to investigate BCR-mediated activation with our different immunogens (**Fig. 4A**). This assay enabled quantification of intracellular calcium signaling, a hallmark of BCR clustering and activation. First, we investigated BCR-mediated activation from the constructs alone to assess the role of micelle formation on B cell activation. Direct incubation of Fluo-8 loaded glVRC01 cells with amph-eOD conjugates significantly increased calcium flux relative to soluble eOD (**Fig. 4B,D**), whereas amph-MD39 constructs showed no difference compared to soluble MD39 (**Fig. 4C,E**). Amph-PEG2K-eOD and amph-PEG5K-eOD variants, which both form micelles of ∼25nm, elicited similar levels of B cell activation that did not significantly differ from each other but were both significantly enhanced from soluble eOD (*p*<0.0001) (**Fig. 4D**). As observed previously, amph-vaccine micelle formation was not influenced by PEG linker length and was instead primarily determined by antigen MW (**Fig. 1**). This is supported by the calcium flux results with amph-MD39, which does not form micelles, as amph-MD39 did not enhance B cell activation over soluble / monovalent MD39 (**Fig. 4E**). Our results indicate that multivalent antigen presentation through micelle formation was responsible for BCR clustering and activation.

**Figure 4.**
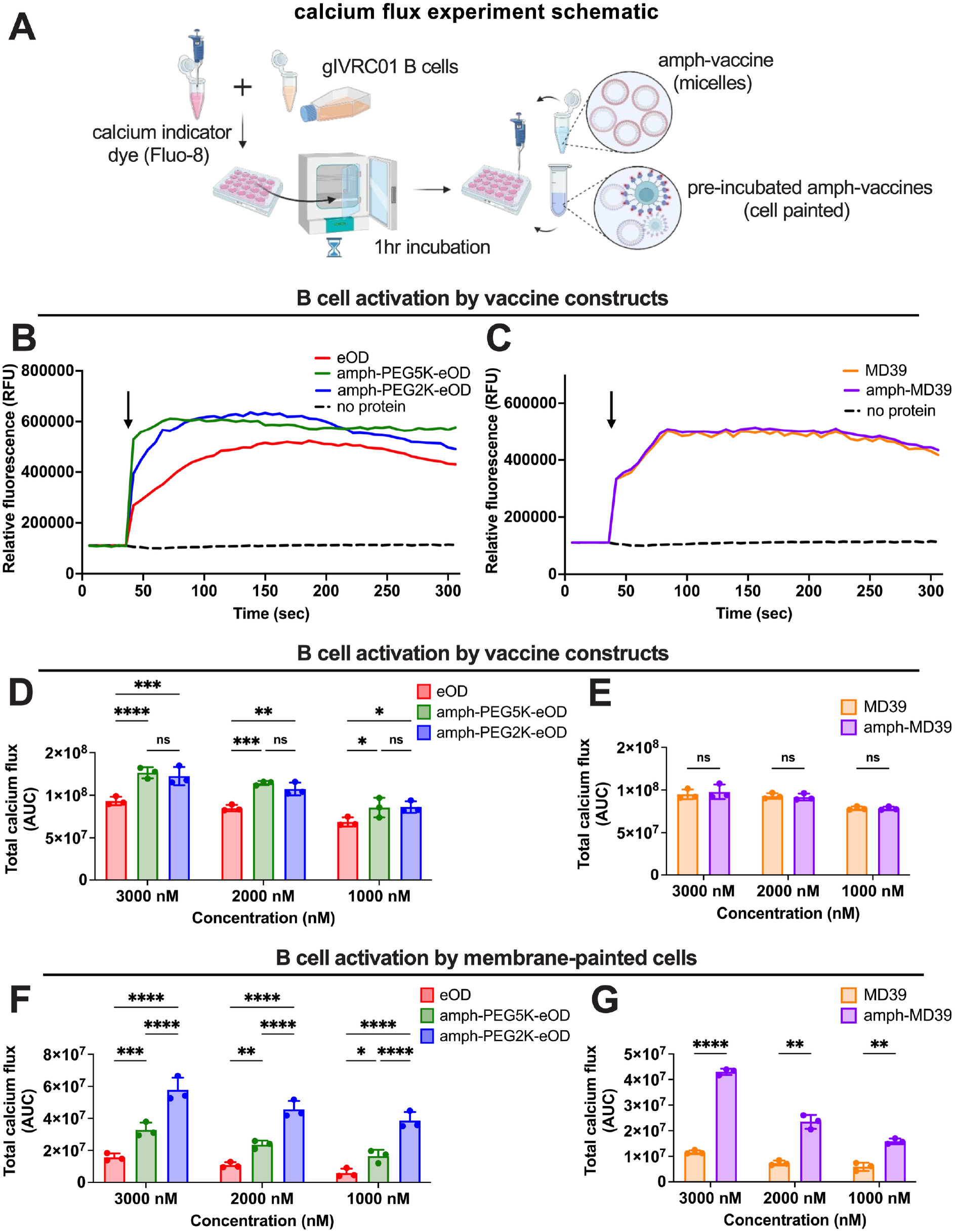
Amphiphile-protein conjugates activate glVRC01 B cells in both micellar and cell painted form. **A)** Experiment schematic for calcium flux assays: gIVRC01 B cells were loaded with calcium indicator (Fluo-8) to monitor calcium flux as a metric of BCR-mediated signaling and activation induced by vaccine immunogen binding. Activation was measured in real-time as Fluo-8 signal upon addition of immunogens (marked by black arrows in (B) and (C)) on a fluorescent plate reader. In a separate independent experiment, Ramos B cells were first pre-incubated with immunogens for 1 hr at 37°C to allow for membrane insertion, then washed Ramos cells were added to dye-loaded glVRC01 cells. **B-C)** Representative calcium flux kinetics for **B)** eOD and **C)** MD39 constructs (3000 nM), showing relative fluorescence units (RFU) over time. **D-E)** Total calcium flux (quantified as integrated area under the curve, AUC, from (B) and (C)) from glVRC01 cell incubation with **D)** eOD and **E)** MD39 *constructs* as a function of antigen concentration. **F-G)** Total calcium flux AUC from glVRC01 cell incubation with **F)** eOD-and **G)** MD39-*painted Ramos cells* as a function of antigen concentration. Statistical significance determined by ordinary one-way ANOVA followed by Tukey’s post hoc test (for eOD groups) or unpaired t-test (for MD39 groups). All data shown are presented as mean ± SEM (n=3).

Next, we investigated BCR-mediated activation from cells pre-incubated and pre-painted with amph-protein to determine the role of membrane insertion on B cell activation. To assess whether amphiphile-mediated cell painting led to multivalent antigen display on nonspecific cells in a manner capable of activating antigen-specific B cells, Ramos cells (not antigen-specific) were pre-incubated with eOD, amph-PEG2K-eOD, amph-PEG5K-eOD, MD39, or amph-MD39 for 1 hour at 37°C to allow for membrane insertion. Then Ramos cells were washed to discard any free (unanchored) antigen and mixed with Fluo-8 loaded glVRC01 cells, and VRC01 cells were measured for calcium flux. Under these conditions, protein amphiphile conjugates significantly enhanced B cell activation compared to soluble unmodified proteins (*p*<0.0001) (**Fig. 4F,G**). In particular, amph-PEG2K-eOD and amph-MD39 each induced three- and four-fold higher calcium flux signaling, respectively, compared to the soluble proteins. Furthermore, amph-PEG2K-eOD outperformed amph-PEG5K-eOD by eliciting two-fold higher response, which tracked with our observations of cell painting. The trends observed here were consistent across all concentrations tested, suggesting a mechanistic behavior whereby membrane insertion of amph-proteins can enhance B cell activation in neighboring antigen-specific B cells **(Fig. S4A-D)**. We hypothesized this might play a role in driving B cell activation in secondary lymphoid organs, so we next evaluated vaccine trafficking to draining lymphatics followed by B cell activation *in vivo*.

### Amphiphile conjugation enhances protein antigen uptake and persistence in draining lymph nodes following subcutaneous injection in mice

Albumin ‘hitchhiking’ promotes lymphatic trafficking by leveraging albumin’s role as a fatty acid transporter, enabling its use as a noncovalent chaperone for LN delivery of lipid-modified cargo (*23–27*). This drug delivery strategy has been demonstrated to enhance LN trafficking of small peptides and molecular adjuvants following subcutaneous injection (*12, 28–30*). We previously showed that DSPE-PEG modification also enhances mucosal uptake and mucosal associated lymphoid tissue trafficking of monomeric proteins following intranasal administration (*1*). We therefore expected that modifying monomeric protein antigens like eOD (those with MW below the threshold expected to passively drain into the lymphatics) with an albumin-binding lipid tail would enhance trafficking into the lymphatics and accumulation in draining LNs after subcutaneous injection. We also hypothesized that cell membrane insertion may lead to enhanced persistence and retention of PEG2K amphiphile vaccines in LNs over time. To test this, we fluorescently labeled soluble or amphiphile-conjugated proteins and evaluated in vivo biodistribution using IVIS following subcutaneous immunization in mice (**Fig. 5A**). Total antigen accumulation in the draining igLNs was quantified by measuring fluorescence (total radiant efficiency) of a defined region of interest at one- and four-days following injection (**Fig. 5B, S5A**). Overall, fluorescence signal was highest at day one post injection and declined for all groups by day four; however, differences were more pronounced between both amph-eOD conjugates and eOD than between amph-MD39 and MD39 at the early timepoint (**Fig. 5C**). At day one, amph-PEG2K-eOD and amph-PEG5K-eOD exhibited approximately 4-to 5-fold greater accumulation in the igLNs compared to soluble eOD (*p*=0.069 for amph-PEG2K-eOD) (**Fig. 5D**), indicating that amphiphile binding to albumin increased drainage into the lymphatics compared to eOD which is more likely to drain into systemic circulation (*51, 52*). Trends were similar at four days post immunization, but differences between amph-PEG2K-eOD and eOD became more pronounced with significantly greater signal retention of amph-PEG2K-eOD compared to eOD (*p*<0.05) (**Fig. 5D**). At this timepoint amph-PEG5K-eOD signal was still elevated from eOD, but did not differ significantly from either eOD or amph-PEG2K-eOD. Given that MD39 has a higher MW (∼77-220 kDa) that falls above the threshold for lymphatic drainage, we did not expect amphiphile modification to significantly alter initial lymphatic trafficking of MD39 at the early timepoint. Indeed, while amph-MD39 was elevated it did not exhibit significantly greater accumulation in igLNs one day after injection compared to MD39, and this difference was smaller than that observed for amph-eOD conjugates (**Fig. 5E,C**). Yet, by four days after injection, amph-MD39 accumulation in igLNs was significantly greater than that of soluble MD39 by 2.5-fold (*p*<0.05). Overall, these data suggest that early-stage lymphatic trafficking was predominantly driven by albumin-binding via the DSPE-lipid tail in amph-eOD vaccines, but our observation that differences in pharmacokinetics became more pronounced over time between PEG2K amphiphile conjugates of both eOD and MD39 compared to their soluble controls suggests that cell painting played a role in promoting persistence of antigen in the dLN at later timepoints.

**Figure 5.**
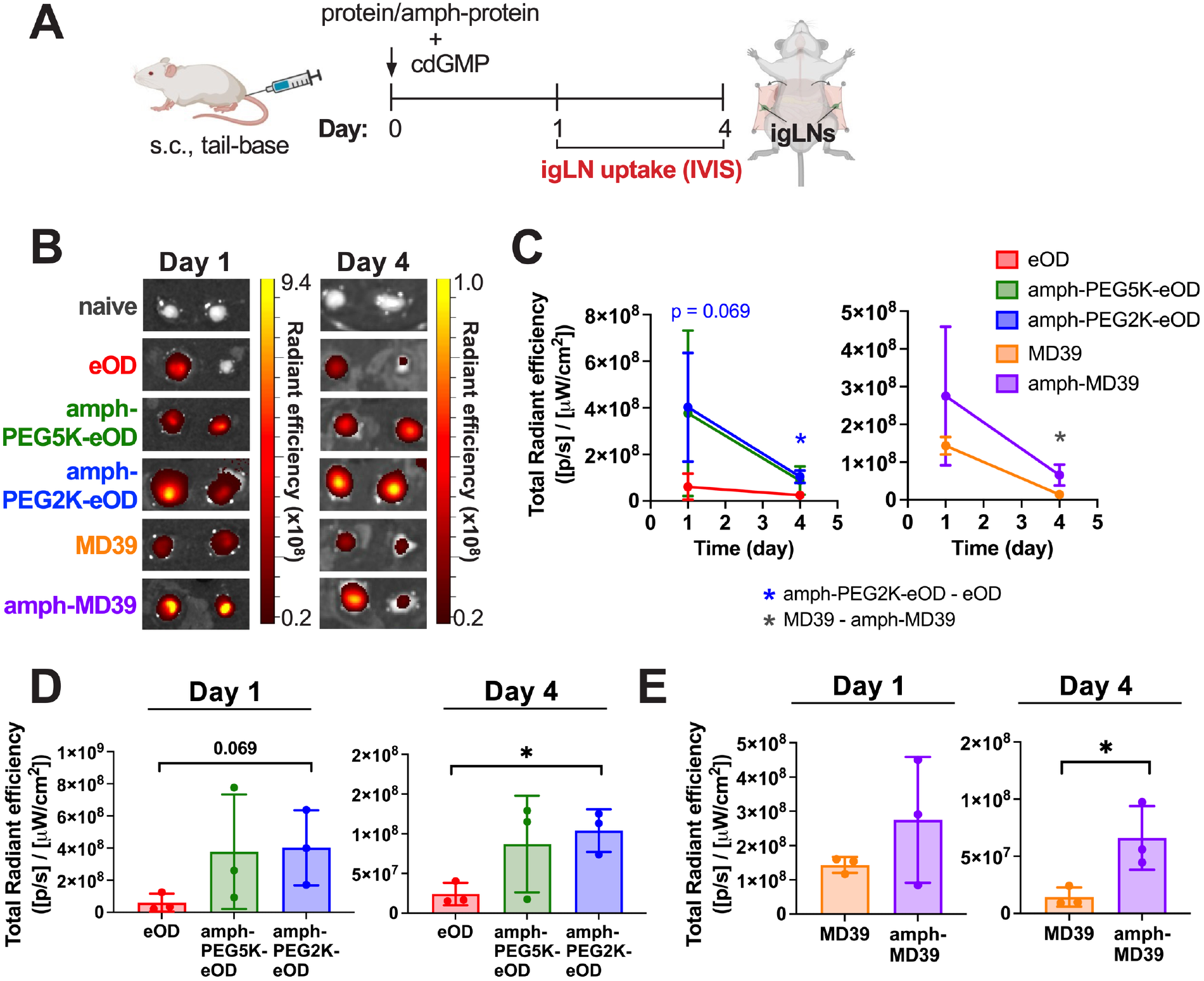
Amphiphile-protein conjugates enhance vaccine uptake and retention in the inguinal lymph nodes. **A)** Experiment schematic: groups of Balb/c mice (n=3 animals per group) were immunized subcutaneously (s.c.) at the tail base with AF647-labeled protein or amph-protein conjugates plus cdGMP adjuvant. Inguinal lymph nodes (igLNs) were isolated and imaged by IVIS at one and four days post-immunization to measure vaccine uptake. **B)** Representative IVIS images of vaccine AF647 signal in igLNs at one and four days post-immunization. **C)** Quantified IVIS signal in igLNs shown as total radiant efficiency over time. Comparison at each timepoint of **D)** eOD groups and **E)** MD39 groups. Statistical significance determined at each time point by ordinary one-way ANOVA followed by Tukey’s post hoc test (for eOD groups) or unpaired t-test (for MD39 groups). All data presented as mean ± SEM.

### Amphiphile conjugation and cell painting enhance germinal center B cell activation in draining lymph nodes *in vivo*

We hypothesized that enhanced vaccine trafficking to and retention in the dLNs would increase antigen availability to B cells and antigen-presenting cells (APCs), while amph-vaccine cell painting could also increase multivalent antigen presentation to B cells, thereby priming a stronger GC response through both mechanisms. To evaluate this, we quantified GC B cells and T follicular helper (Tfh) cells in the draining igLNs 12 days after subcutaneous immunization in mice (**Fig 6A**). GC B cells were defined as B220⁺CD38⁻GL7^+^ cells gated on live CD3e^-^ single cells (**Fig. S6A)**; Tfh cells were defined as CD4⁺CD44^+^PD-1^+^CXCR5⁺ cells gated on live B220^-^ single cells (**Fig. S7A**). Antigen-specific GC B cells were defined as GC B cells double positive for two fluorescently labeled protein antigen tetramers (fluorophore-streptavidin coupled to biotinylated proteins): RB613-eOD/MD39 and RB780-eOD/MD39. In parallel, early T cell activation was evaluated by measuring the inducible costimulatory response (ICOS) on CD4⁺CD44⁺ T cells. For both eOD and MD39 proteins, amphiphile conjugation significantly enhanced total and antigen-specific GC B cell responses over soluble proteins (*p*<0.0001, **Fig. 6B,C**). Immunization with amph-PEG2K-eOD resulted in nearly five-fold increase in total GC B cells and four-fold increase in antigen-specific GC B cells compared to soluble eOD (**Fig. 6B**), while amph-MD39 produced a 1.5-fold increase in total GC B cells and two-fold increase in antigen-specific GC B cells compared to soluble MD39 (**Fig. 6C**). Notably, amph-PEG2K-eOD also induced significantly greater total GC B cell response than amph-PEG5K-eOD (*p*<0.0001), indicating that cell painting with PEG2K conjugate influenced the overall magnitude of GC B cell activation. Analysis of CD4⁺ T cell subsets revealed similar trends. Both amph-eOD conjugates induced significantly greater Tfh cell expansion than eOD (*p*<0.0001), and amph-MD39 likewise induced greater Tfh cells than MD39 (*p*<0.01) (**Fig. 6D**). Immunization with amph-PEG2K-eOD induced a six-fold greater PD-1⁺ Tfh cell response and eight-fold greater ICOS⁺CD4⁺CD44^+^ T cell response compared to soluble eOD (**Fig. S7C**), demonstrating both enhanced Tfh differentiation and T cell activation. Amph-PEG2K-eOD also induced significantly more Tfh cells than amph-PEG5K-eOD (*p*<0.0001), indicating that cell painting plays a role in promoting Tfh activation and differentiation. Of note, the magnitude of total and antigen-specific GC B cells induced by amph-MD39 immunization were roughly two- and three-fold that of amph-PEG2K-eOD immunization, respectively, while the magnitude of Tfh cells were similar (**Fig. 6B,C**). However, for antigen-specific GC B cells this difference could be partly attributed to the different tetramers used (eOD versus MD39), given that trimeric MD39 has higher valency and avidity than eOD, and thus an MD39-tetramer would be expected to bind more lower affinity B cells than an equivalent eOD-tetramer.

**Figure 6.**
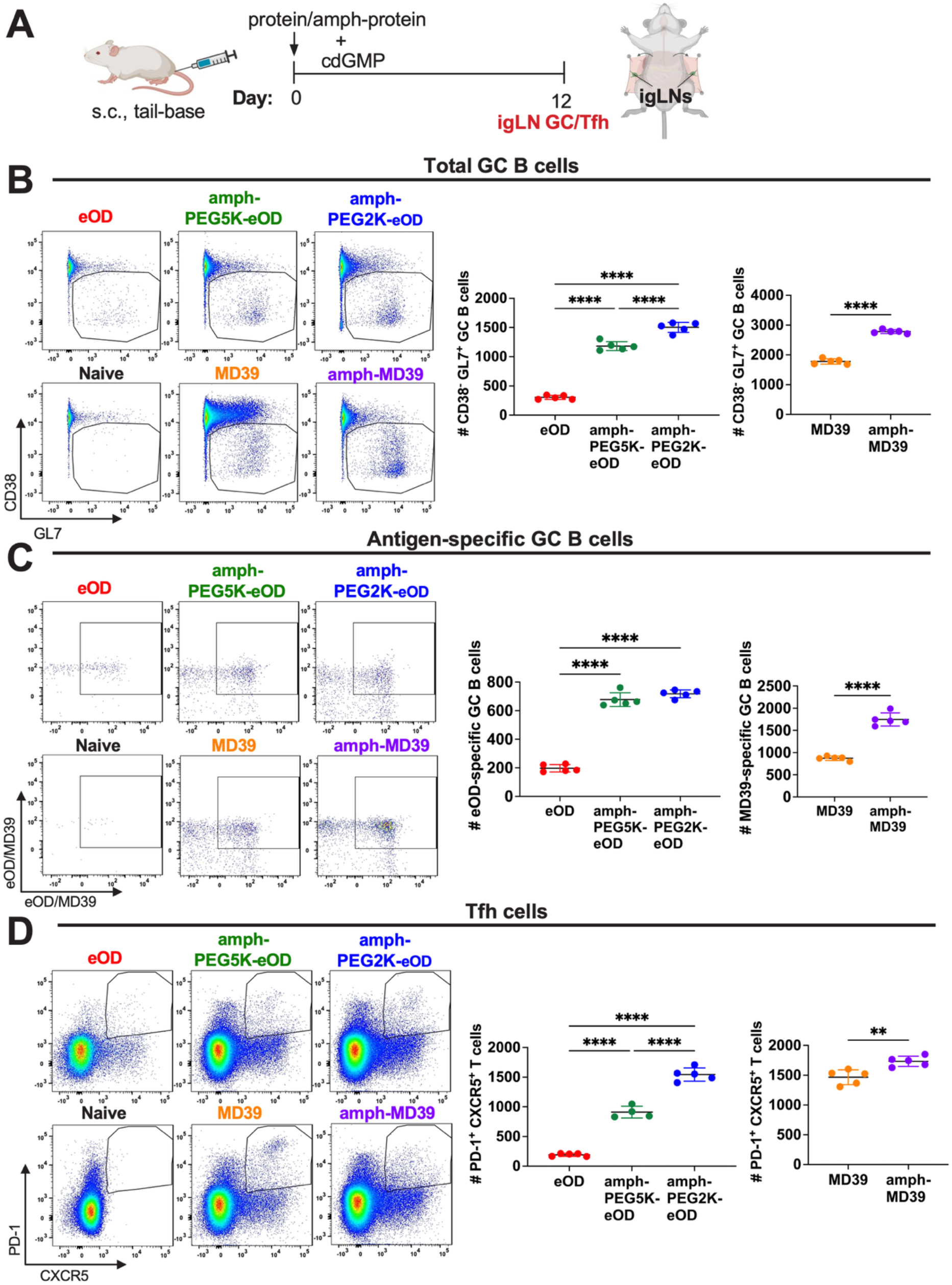
Albumin-binding amphiphile-protein conjugates enhance germinal center and Tfh cell responses. A) Experiment schematic: groups of Balb/c mice (n=5 animals per group) were immunized subcutaneously (s.c.) at the tail base with protein or amph-protein conjugates plus cdGMP adjuvant. Inguinal lymph nodes (igLNs) were isolated on day 12 post-immunization for GC B cell and Tfh cell analysis by flow cytometry. **B-D)** Representative flow cytometry gating and enumeration of **B)** total GC B cells, **C)** antigen-specific GC B cells, and **D)** Tfh cells. Statistical significance determined by ordinary one-way ANOVA followed by Tukey’s post hoc test (for eOD groups) or unpaired t-test (for MD39 groups). All data presented as mean ± SEM. \**p*<0.05, \*\**p*<0.01, \*\*\**p*<0.001, and \*\*\*\**p*<0.0001.

### Amphiphile-protein vaccines induce long-lasting systemic antibody responses following subcutaneous immunization in mice

We next evaluated serum antibody responses elicited by subcutaneous immunization with either soluble or amphiphile-conjugated protein antigens. The magnitude and persistence of serum antibody titers are closely correlated with vaccine efficacy and are often employed as surrogate markers of protective immunity in both preclinical and clinical studies (*53*). Antigen-specific IgG titers were quantified longitudinally by ELISA at biweekly intervals to capture the magnitude and durability of the humoral immune response (**Fig. 7A**). Amphiphile conjugation of both eOD and MD39 significantly enhanced antigen-specific serum IgG compared to soluble protein over 12 weeks (**Fig. 7B,C**). Of note, amph-PEG2K-eOD and amph-PEG5K-eOD both induced high serum IgG titers post-boost that significantly exceeded soluble eOD (*p*<0.001 and *p*<0.0001), but amph-PEG2K-eOD also induced high titers at early timepoints as early as wk2 that significantly exceeded both eOD (*p*<0.01, *p*<0.001) and amph-PEG5K-eOD (*p*<0.0001) pre-boost, indicating strong early priming of the humoral response from cell painting (**Fig. 7B**). Post-boost, amph-PEG-2K and amph-PEG5K-eOD immunization both sustained high antigen-specific IgG titers of around 10^6^-10^7^ through week 12, more than 100-fold greater than soluble eOD, but amph-PEG2K-eOD remained significantly elevated over amph-PEG5K-eOD at all timepoints except week 8. Similarly, amph-MD39 induced significantly higher titers than soluble MD39 both pre-boost at wk4 and post-boost starting at wk8 (*p*<0.0001), roughly 50-to 100-fold greater than MD39 (**Fig. 7C**). These results highlight the broad applicability of amphiphile modification to enhance humoral immunogenicity, even for large multimeric antigens like MD39, while also suggesting that cell painting with PEG2K amphiphile conjugates further enhances humoral activation over PEG5K amphiphile conjugates.

**Figure 7.**
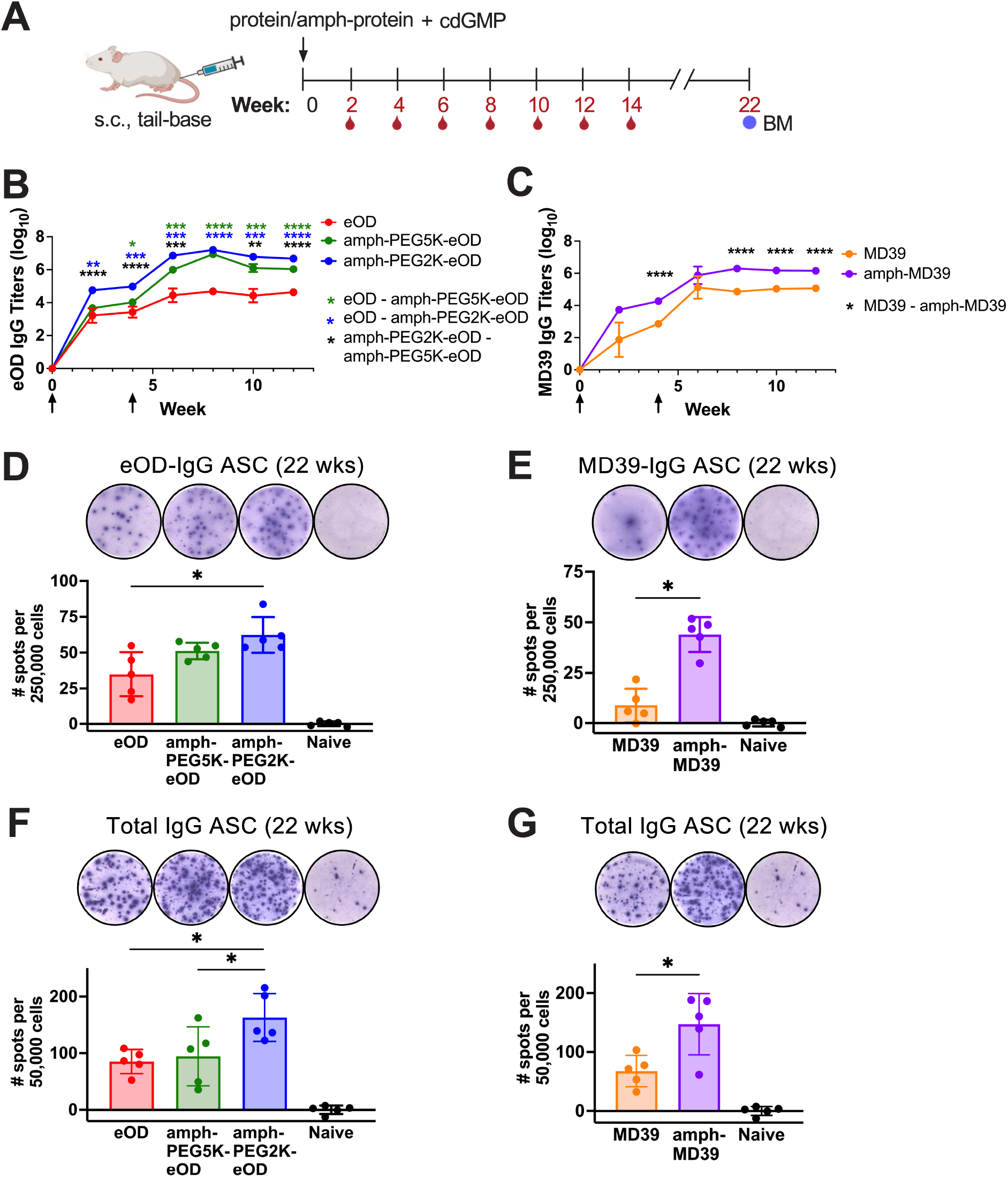
Albumin-binding amphiphiles elicit enhanced systemic immune responses and long-lived bone marrow plasma cell responses. **A)** Experiment schematic: groups of BALB/c mice (n = 5 animals per group) were immunized subcutaneously (s.c.) at the tail base with protein or amph-protein conjugates + cdGMP and boosted 4 weeks later with the same formulations. Serum samples were collected biweekly through week 12 for serum IgG analysis by ELISA, and bone marrow (BM) was isolated at week 22 for antibody-secreting cell (ASC) analysis by ELISPOT. **B-C)** Antigen-specific serum IgG titers measured over time for **B)** eOD groups and **C)** MD39 groups, where black arrows indicate vaccination timepoints. **D-G)** BM IgG ASCs assessed by ELISPOT after 22 weeks, showing **D)** eOD-specific IgG ASCs, **E)** MD39-specific IgG ASCs, **F)** total IgG ASCs for eOD groups, and **G)** total IgG ASCs for MD39 groups. Statistical significance in (B) and (C) was determined by ordinary two-way ANOVA followed by Holm-Sidak’s post hoc test (for eOD groups) or Sidak’s post hoc test (for MD39 groups). Statistical significance in (D) to (G) was determined by ordinary one-way ANOVA followed by Tukey’s post-hoc test (for eOD groups) or unpaired t-test (for MD39 groups). All data presented as mean ± SEM. \**p*<0.05, \*\**p*<0.01, \*\*\**p*<0.001, and \*\*\*\**p*<0.0001.

To determine if high serum antibody responses were accompanied by generation of long-lived plasma cells (LLPC), a metric of durability of the immune response, we then performed ELISPOT assays on bone marrow harvested at the study endpoint (week 22). LLPCs that migrate to the bone marrow niche are the primary source of durable, high-affinity IgG and serve as a hallmark of a successful GC reaction (*7, 14*). Consistent with our GC and serum ELISA data, mice immunized with amph-PEG2K conjugates of eOD and MD39 demonstrated significantly greater frequency of total and antigen-specific IgG antibody-secreting cells (ASCs) in the bone marrow compared to soluble proteins (*p*<0.05) (**Fig. 7D-G**). Amph-MD39 in particular exhibited almost five-fold increase in antigen-specific IgG ASCs compared to soluble MD39 (**Fig. 7E**). The magnitude of ASCs between eOD and MD39 immunogen groups was similar, with amph-PEG2K-eOD and amph-MD39 inducing ∼150 total IgG ASCs per 50,000 cells, roughly two-fold greater than soluble protein (**Fig. 7F,G**). Furthermore, immunization with amph-PEG2K-eOD also induced more total, but not antigen-specific, IgG ASCs in bone marrow than amph-PEG5K-eOD (*p*<0.05) (**Fig. 7F,D**). Interestingly, this trend aligns with our observation that amph-PEG2K-eOD enhanced total, but not antigen-specific, GC B cell responses in igLNs compared to amph-PEG5K-eOD (**Fig. 6B,C**). It might suggest that multivalent antigen presentation through cell painting with amph-PEG2K-eOD drives more polyclonal B cell activation and expansion on top of eOD-specific clonal expansion compared to amph-PEG5K-eOD. Overall, these data support the hypothesis that enabling cell painting through amphiphilic modification of protein antigens with a short PEG linker is a key driver of sustained humoral immunity.

## DISCUSSION

We previously demonstrated that modifying protein antigens with amphiphilic lipid tails promotes albumin-mediated uptake across mucosal barriers following intranasal administration, leading to enhanced B cell priming and systemic and mucosal humoral immune responses (*1*). Other work has demonstrated that amphiphile conjugates of peptide antigens, haptens, and CpG molecular adjuvants exhibit enhanced lymphatic trafficking and antigen retention in LNs following subcutaneous administration, leading to enhanced T cell priming and cellular immune responses (*12, 28, 32*). Furthermore, we and others have demonstrated the ability of amphiphile conjugates to insert into cell membranes (*1, 28, 32*). Zhang et. al. observed membrane insertion with amph-peptide and -hapten conjugates, and Liu et. al. with amph-nucleotide conjugates (*12, 32*). *However, membrane insertion by amphiphile conjugates has not previously been studied in the context of B cell activation with protein immunogens, nor was the interplay between molecular properties such as antigen molecular weight, PEG linker length, and molecular conformation fully understood.* Here, we demonstrate that membrane insertion, aka ‘cell painting’, is a key behavior that plays a significant role in driving B cell activation and humoral immune responses from amphiphile-protein conjugate vaccines. To our knowledge we are the first to demonstrate this mechanism with amphiphile-protein conjugates and investigate it as a strategy for B cell activation. By comparing amphiphile conjugates with varying antigen sizes and PEG linker lengths, we identified molecular design parameters that govern the transition between amphiphile micelle formation and cell painting, then investigated how these distinct conformational states influenced the magnitude of B cell activation and subsequent immune responses.

Our *in vitro* characterization studies first elucidated how conformational behavior of amph-vaccines was dependent on molecular properties such as PEG length and protein MW. Membrane insertion was driven by PEG linker length (favored with shorter PEG linkers), while micelle formation was driven by protein MW (sterically hindered with larger trimer protein). Yet, we observed that all amphiphile conjugates (amph-PEG2K-eOD, amph-PEG5K-eOD, and amph-MD39) bound albumin *independent* of antigen MW or PEG linker length. We predicted this would enable albumin hitchhiking for enhanced LN trafficking *in vivo* for smaller MW proteins such as eOD, but would not significantly alter early LN trafficking patterns for larger proteins such as MD39. This is because lymphatic trafficking is known to function in a MW-dependent fashion based on the size-selective permeability of the blood vascular endothelium (*47, 54*). While small unmodified proteins like eOD are rapidly cleared into the systemic circulation, amphiphile conjugation allows them to bind albumin (∼67 kDa) as a chaperone, shifting the effective MW of the vaccine complex above the threshold for effective passage into lymphatics (*1, 47, 55*). While we did observe these trends in our IVIS trafficking data at day 1, it was not statistically significant. However, for both eOD and MD39 antigens, amph-PEG2K conjugation significantly enhanced *persistence* in LNs by day 4. Of note, LN persistence was significantly enhanced with amph-PEG2K-eOD but not amph-PEG5K-eOD, suggesting another mechanism at play besides albumin hitchhiking. We did not attribute enhanced persistence (observed with both eOD and MD39 conjugates) to micelle formation, because while amphiphile conjugates of monomeric eOD formed stable micelles, larger MD39 trimer conjugates were sterically hindered from micelle formation. Yet both amph-PEG2K-eOD and amph-MD39 conjugates demonstrated stable cell membrane insertion. LN persistence was thus attributed to cell painting as the common property shared between amph-PEG2K-eOD and amph-MD39.

Combined, our *in vitro* and *in vivo* results demonstrated that membrane insertion, rather than micelle assembly, was the primary driver of B cell activation with amph-protein vaccines – both *in vitro* in human glVRC01 cells and *in vivo* in mouse GC B cells. Optimizing PEG linker length allowed for tunable control over cell painting behavior, significantly reduced with longer PEG5K. Cell painting with amph-PEG2K conjugates transformed the cell surface into a multivalent display of antigen, providing the BCR crosslinking necessary for potent B cell activation as shown *in vitro* by fluorescence microscopy and calcium flux assays. Our *in vivo* results subsequently showed significant activation of GC B cells and Tfh cells in dLNs along with sustained serum antigen-specific IgG antibody responses from amph-PEG2K-eOD and amph-MD39 over amph-PEG5K-eOD / eOD and MD39, respectively. This aligned with a significant increase in the long-lived antibody-secreting plasma cells (ASCs) within the bone marrow for amph-PEG2K-eOD and amph-MD39 compared to other groups, findings that suggest cell-painting can effectively prime a larger pool of high-affinity B cells that differentiate into long-lived plasma cells (LLPCs) in BM – of clinical importance since LLPCs serve as the primary source of sustained, high-avidity protective antibodies for extended periods post-immunization (*56*). Of note, while significant differences were observed between amph-PEG2K-eOD and amph-PEG5K-eOD across all other humoral metrics, they did not differ significantly for *antigen-specific* GC B cells or *antigen-specific* ASC responses. This suggests that amph-eOD cell painting with PEG2K conjugates may drive polyclonal activation over monoclonal activation. Overall, these findings reveal that cell painting by amphiphile vaccines provides a superior platform for B cell engagement and activation in GCs, leading to significantly higher antibody titers and long-lived plasma cell generation compared to conventional soluble protein-based vaccines or amphiphile-protein vaccines do not exhibit cell painting.

B cell activation and GC responses are believed to be a critical precursor of an effective antibody response for many infectious diseases, in particular highly variable pathogens like HIV. GCs serve as a training ground for a high affinity antibody response; vaccine efficacy largely relies on a vaccine’s ability to produce this training ground. Thus, enhancing B cell and GC activation through a mechanism of a vaccine platform that’s adaptable to different types of antigens, ranging from monomers to multimers, is likely to be of interest in developing vaccines for diverse pathogens. We know antigen availability in LNs is a major driver of the GC response (*55*). While soluble antigens tend to be cleared via the subcapsular sinus, cell painting appears to tether amphiphile conjugates to local cells which may prolong antigen availability to cognate B cells in the GC. This membrane-anchored state may also provide follicular protection by partially shielding the protein’s antigenic epitopes from extracellular digestion by resident proteases (*57, 58*). Preservation of conformational epitopes is particularly crucial for structurally sensitive immunogens like MD39, where the maintenance of the native-like trimer state is necessary for eliciting broadly neutralizing antibodies (*44, 59–61*). In addition, our studies indicate that membrane-painted cells are capable of multivalently presenting antigen to nearby B cells, priming the GC response through enhanced B cell activation.

Our studies focused on two protein antigens derived from HIV viral envelope, but the implications of the cell painting mechanism are applicable across disease settings. While we utilized eOD and MD39 as models for small monomer and large multimer protein antigens, respectively, our results established that the amph-vaccine platform is well suited for antigens with a broad range of MWs. We have shown that immunogenicity of both small and structurally large immunogens can be enhanced through amphiphile-conjugation and cell painting. This is likely to be of value in developing vaccines for diverse pathogens as antigen targets can differ greatly across disease settings. Depending on pathogen, an optimal antigen target may consist of a large conformationally-dependent trimer like MD39 or the highly glycosylated hemagglutinin (HA) protein from influenza (*62*), rather than a 20 kDa monomer like eOD or the receptor binding domain (RBD) (*1*) from SARS-CoV-2. Ultimately, the findings of this study support cell painting with amph-protein vaccines as a promising strategy to enhance humoral immunity and subunit vaccine efficacy.

This work builds upon previous studies of cell membrane insertion with haptens and peptide antigens (*28, 32*), here expanded to different MW protein antigens while identifying cell painting as a mechanism for B cell activation. Our results show enhanced membrane insertion with amph-PEG2K-eOD compared to amph-PEG5K-eOD, corroborating observations by Zhang et. al. that showed amphiphile cell membrane insertion efficiency was dependent on PEG linker length with longer PEG linkers destabilizing membrane insertion (*32*). Previous studies by Liu et. al. and Yu et. al. also highlighted that amphiphiles with short polymer chains were rapidly inserted into the cell membrane. They observed that while insertion stability increased with *shorter* PEG, it also increased with *longer* lipid, concluding that chain length of both polymer and lipid impacts membrane insertion state (*12, 63*). For example, Liu, Kwong, and Irvine demonstrated that diacyl lipid conjugates of oligonucleotides exhibited significantly greater membrane insertion than conjugates using cholesterol or single chain C18 forms of lipid (*50*). These studies demonstrated that choice of lipid in the platform, in addition to PEG linker length, is an important consideration for achieving a high degree of cell painting.

Lipid-mediated membrane anchoring is not a new phenomenon, of course; it occurs endogenously in cell biology processes as well as with existing drugs. For example, it mirrors the natural biological processes of myristoylation and palmitoylation, where cells utilize fatty acids to anchor signaling proteins into specific domains within the cell membrane to facilitate high-avidity molecular interactions (*64*). Many existing drugs take advantage of analogous physicochemical behavior to extend half-life *in vivo*. For example, peptides with fatty acid side chains, like insulin detemir (‘Levemir’, containing a C14 myristoyl side chain), insulin degludec (‘Tresiba’, containing a C16 hexadecanedioic acid side chain), and liraglutide (GLP-1 receptor agonist containing a C16 palmitic acid side chain), were all designed with a lipid tail to bind serum albumin to extend circulation times (*65–67*). As albumin binding occurs reversibly, amphiphilic drugs also hold the potential to insert into cell membranes, leading to altered pharmacokinetics through tissue persistence. Established antibiotic drugs like the lipopeptide daptomycin (*68*) and lipoglycopeptides (dalbavancin, oritavancin, telavancin) (*69*) use acyl tails to anchor into bacterial cell membranes. Cationic amphiphilic drugs such as the antiarrhythmic drug amiodarone (*70*) also partition into cell membranes, where they can induce a disorder characterized by excessive cellular phospholipid accumulation known as phospholipidosis. For drugs with B cell epitopes, our results suggest that membrane insertion may lead to immune activation through multivalent B cell activation, which could accelerate anti-drug antibody responses. Thus, this work raises important questions about the potential of cell painting to generate unappreciated immune effects with amphiphilic drugs.

One limitation of this present study is the challenge of translating immunological results from small animal models to humans. Despite this limitation, it’s worth noting that our *in vivo* results in mice corroborated with our *in vitro* results in human cell lines. Our results in mice demonstrated a clear link between a vaccine’s cell-painting efficiency and its ability to activate robust GC responses, stimulate high IgG titers, and generate long-lasting ASCs in the bone marrow *in vivo*. This structure-function relationship between vaccine properties, cell painting, and B cell activation was similarly observed *in vitro* in Ramos and glVRC01 human cell lines. Given that increased durability of GC reactions is essential for triggering superior functional immunity in humans, a strategy that enhances B cell and GC activation is likely to be translatable across models (*71, 72*). However, while we identified cell painting as a driving mechanism of humoral activation, we did not identify which specific cells *in vivo* were involved in cell uptake and membrane insertion with this work. Yet we touched on this in our previous work, where we demonstrated that amph-vaccines exhibited enhanced uptake in macrophages, dendritic cells, and B cells (*1*). Another remaining question is whether choice of accompanying adjuvant affects membrane insertion behavior. The present studies were performed using cdGMP, a cyclic dinucleotide small molecule adjuvant. If a lipophilic or amphiphilic adjuvant were used instead – such as iscomatrix or saponin MPLA nanoparticles (SMNP) – would those molecules affect membrane insertion through competitive association with the cell membrane or lipophilic self-association? These specific nuances of cell painting remain to be investigated in future work.

While our work here focused on parenteral immunization (subcutaneous injection) to avoid convoluting variables from mucosal uptake, the amphiphile platform holds significant promise for mucosal administration. Mucosal vaccination is attractive for generating mucosal IgA and resident memory cells as frontline defenses at mucosal barrier tissues where transmission takes place (*73, 74*). Our previous work demonstrated that intranasal amphiphile-protein vaccines effectively crossed mucosal barriers in the nose via albumin-mediated FcRn transcytosis, enhancing GC activation in nasal associated lymphoid tissue (NALT) along with systemic and mucosal antibody responses (*1*). We hypothesize that cell painting may play an important role for amph-vaccines following mucosal vaccination, possibly an even greater role than that in parenteral administration, by enhancing retention, reducing clearance, and aiding uptake in mucosal epithelium. Thus, ongoing and future work in our lab will explore the role of cell painting for enhancing mucosal humoral immune activation with intranasal amph-vaccines.

To conclude, this work established that amphiphile conjugation of protein antigens provides a modular platform for enhancing humoral immune responses, driven in part by a mechanism of cell painting. Using an engineered amphiphile vaccine platform, we investigated three key design mechanisms: self-assembly into nanoscale micelles, noncovalent albumin binding for improved lymphatic transport, and cell membrane insertion (cell painting) for multivalent antigen display and B cell activation. Our findings strongly suggest that cell painting by amphiphile-conjugated proteins both enhance persistence of albumin-hitchhiking vaccines in dLNs while also achieving multivalent B cell activation, serving as a key driver of humoral immunity. Modulating PEG linker length allowed for control over cell painting, while both small monomeric and large trimer antigens benefited from cell painting. These results provide evidence for a mechanism behind the superior immunogenicity of amphiphile-protein vaccines, secondary to albumin hitchhiking, and suggest that cell painting can be incorporated into vaccine design to enhance efficacy and humoral immunity.

## MATERIALS AND METHODS

### Study design

The major objective of this study was to evaluate molecular and cellular mechanisms of amphiphile-protein conjugate vaccines, and in particular identify the contribution of membrane insertion (cell painting) versus micelle formation on vaccine efficacy for B cell activation and humoral immunity. Our hypothesis was that membrane insertion from amph-vaccines plays a role in promoting humoral vaccine efficacy by increasing multivalent B cell activation and enhancing persistence of vaccine in dLNs, so our studies were designed to evaluate B cell activation in vitro and in vivo with different conformations of amph-protein vaccines. To evaluate the role of membrane insertion, we compared soluble and amphiphile forms of clinically relevant HIV-env model antigens (eOD gp120 monomer and MD39 SOSIP trimer), including amphiphiles with varying PEG linker lengths resulting in different conformational and membrane insertion properties: eOD, amph-PEG2K-eOD, amph-PEG5K-eOD, MD39, amph-PEG2K-MD39. Amph-PEG5K-eOD was included as a control for cell painting, as it exhibits equivalent albumin binding and micelle formation but significantly reduced membrane insertion compared to amph-PEG2K-eOD. Amph-MD39 was included as a control for micelle formation, as it exhibits albumin binding and membrane insertion but no micelle formation. We first quantified membrane insertion or ‘cell painting’ in vitro in nonspecific cells, and investigated whether painted cells could induce functional BCR-mediated activation in vitro in antigen-specific glVRC01 B cells by comparing the B cell activation from soluble antigens, amph-antigen micelles, or amph-antigen painted cells. Mice were then immunized with protein or amph-protein conjugates plus cyclic di-GMP (cdGMP) adjuvant via subcutaneous tail base injection to assess early and local vaccine kinetics and immune activation in vivo as a function of cell painting, as well as humoral immunity over time. Vaccine trafficking to and persistence in dLNs 1- and 4-days post immunization was assessed by IVIS. Germinal center (GC) B cell and T follicular helper (Tfh) cell activation 12 days post immunization was assessed by flow cytometry. Longitudinal serum antibody responses and long-lasting plasma cells were assessed by ELISA and ELISPOT, respectively. For exclusion criteria, flow cytometry data were omitted if the sample’s total cell counts were less than 3,000.

### eOD and MD39 protein production

HIV Env gp120 engineered outer domain (eOD) protein with N-terminal cysteine and C-terminal PADRE universal helper T cell epitope (AKFVAAWTLKAAA) was produced in Expi293F human embryonic kidney (HEK) cells (Thermo Fisher Scientific), as previously described (*13, 44, 59*). HIV BG505 SOSIP MD39 trimer containing a C-terminal cysteine was also expressed in Expi293F HEK cells, as previously described (*59, 61*). Proteins were purified on a nickel affinity column, followed by the removal of free salts using a Zeba spin desalting column with a 7 kDa molecular weight cutoff (MWCO) (Thermo Fisher Scientific). Anti-human VRC01 (hVRC01) antibody was synthesized and generously provided by Darrell Irvine.

### Amph-PEG2K-eOD and amph-PEG5K-eOD conjugation and purification

Amph-PEG2K-eOD was synthesized and purified as previously described (*1*). Briefly, eOD protein (≥1 mg/ml) was first reduced using 10 molar equivalents of tris(2-carboxyethyl)phosphine (TCEP) for 15 minutes at 25°C. Excess TCEP was removed by centrifugal filtration with 10 kDa MWCO Amicon spin filters, washing the protein three times with phosphate-buffered saline (PBS). The reduced protein (1 to 5 mg/ml) was then reacted with 4 molar equivalents of dried DSPE-PEG2K-maleimide (1,2-distearoyl-sn-glycero-3-phosphoethanolamine-N-[maleimide(polyethylene glycol)-2000]) or DSPE-PEG5K-maleimide (1,2-distearoyl-sn-glycero-3-phosphoethanolamine-N-[maleimide(polyethylene glycol)-5000]) (Avanti Polar Lipids) in PBS for 2 hours at 25°C with intermittent vortexing, followed by gentle mixing for 18 hours at 4°C. Amphiphile conjugates were purified by size exclusion chromatography (SEC) using a Sepharose CL6B gravity column (Sigma-Aldrich) eluted with PBS. Intrinsic tryptophan fluorescence (excitation 280 nm, emission 340 nm) was used to detect fractions containing conjugated protein-amphiphile micelle versus unconjugated protein peaks using a multimodal plate reader (SpectraMax iD3, Molecular Devices, San Jose, CA, USA). Micelle peak fractions were collected, concentrated using 10 kDa MWCO Amicon spin filters, and quantified by UV-Vis spectrophotometry (Nanodrop One, Thermo Fisher Scientific). Amphiphile concentration was determined by quantifying the protein peak at 280 nm, corrected for lipid scattering by subtracting the background lipid absorbance power function fitted from 310-500 nm. Particle size was analyzed by dynamic light scattering (DLS) (Zetasizer Nano, Malvern) following dilution in PBS.

### Amph-MD39 conjugation and purification

MD39 trimer (≥1 mg/ml) was reduced with 10 molar equivalents of tris(2-carboxyethyl)phosphine (TCEP) for 15 minutes at 25°C. TCEP was removed by centrifugal filtration using 10 kDa MWCO Amicon spin filters with three washes of phosphate-buffered saline (PBS). The protein (1 to 5 mg/ml) was then reacted with 5 molar equivalents of DBCO-PEG4-maleimide (dibenzocyclooctyne-PEG4-maleimide) in PBS for 18 hours at 4°C. Unreacted maleimide-PEG4-DBCO was removed using 10 kDa MWCO Amicon spin filters. The product was analyzed by UV-Vis spectrophotometry (Nanodrop One, Thermo Fisher Scientific) for a DBCO peak at 309 nm. Next, MD39-DBCO was mixed (≥1 mg/ml) with 5 molar equivalents of dried DSPE-PEG2K-azide (1,2-distearoyl-sn-glycero-3-phosphoethanolamine-N-[azido(polyethylene glycol)-2000]) in PBS for 2 hours at 25°C with intermittent vortexing, followed by gentle mixing for 18 hours at 4°C. The reaction’s completion was verified by UV-Vis with the absence of the DBCO peak at 309 nm. MD39 concentration was determined by quantifying the protein peak at 280 nm, corrected for lipid scattering as described above. Particle size was analyzed by dynamic light scattering (DLS) following dilution in PBS.

### Fluorophore labeling

For IVIS and membrane insertion studies, vaccine antigens eOD, amph-PEG2K-eOD, amph-PEG5K-eOD, MD39, and amph-MD39 were labeled with AlexaFluor647 (AF647) N-hydroxysuccinimide (NHS) ester (Thermo Fisher Scientific) by reacting the fluorophore with the soluble protein or amphiphile conjugate (≥1 mg/ml) in 0.1 M sodium bicarbonate buffer for 1 hour at 25°C, per the manufacturer instructions. hVRC01 was labeled with AlexaFluor555 (AF555) N-hydroxysuccinimide (NHS) ester (Thermo Fisher Scientific) by reacting the fluorophore with hVRC01 (≥1 mg/ml) in 0.1 M sodium bicarbonate buffer for 1 hour at 25°C, per the manufacturer instructions. Labeled proteins were purified using 10 kDa MWCO Amicon spin filters, and the degree of labeling (DOL) was quantified by UV-Vis.

### Micelle morphology by transmission electron microscopy (TEM)

Protein and amphiphile-protein conjugates were diluted to a concentration of 500 nM in PBS and vortex-mixed for 30 seconds prior to application. The surface of a 400-mesh ultrathin carbon film on a lacey carbon-supported copper grid (Ted Pella, Redding, CA) was negatively charged by glow discharge for 30 seconds. A 3 µL aliquot of each sample was deposited onto the grid for 30 seconds at 22°C. Excess liquid was carefully blotted using Whatman® filter paper (Schleicher & Schuell, Germany). The stained grids were then imaged using a FEI G2 F30 S-TWIN transmission electron microscope (FEI, Hillsboro, OR) operated at 300 kV. Micelle diameters were calculated post-imaging using Fiji software.

### Albumin binding

To further characterize and quantify albumin binding, fatty acid-free BSA was covalently immobilized on NHS-activated agarose resin. First, 20 mg of fatty acid-free BSA was dissolved in 3 mL PBS and added to 250 mg NHS-activated agarose resin (Pierce™ NHS-Activated Agarose, Thermo Scientific). The coupling reaction was allowed to proceed for 2 hours at 25°C. Unbound BSA was removed by centrifugation, and the resin was quenched using 50 mM Tris-HCl buffer (pH 8.0). The beads were washed extensively with PBS to remove residual reactants. AF647-labeled soluble protein or amph-protein was then mixed with 300 µL of the BSA-agarose beads (50 mg dry resin/mL) at a final antigen concentration of 600 nM in PBS. The mixtures were incubated at 37 °C for 1 hour with gentle agitation. After binding, beads were separated from the filtrate buffer using Pierce^TM^ spin columns (Cat# 89896), and AF647 fluorescence measurements were made using a Bio-Rad Chemidoc gel imager (ex/em: 647 nm / 670 nm). Bound protein was visualized and quantified by evaluating the fluorescence intensity using ImageJ. Binding efficiency was calculated as:

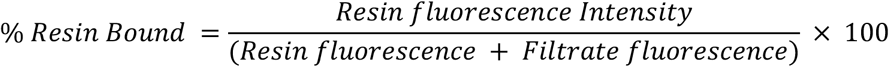

To evaluate the percentage of albumin-binding at different BSA concentrations, we performed a co-elution assay using size exclusion chromatography (SEC) and fluorescence imaging. AF647-labeled soluble protein or amph-protein at 1500 nM was incubated with varying concentrations of AF488-labeled BSA ranging from 250 - 6500 nM in PBS (pH 7.4) at 37 °C for 1 hour. The mixture was separated by SEC using Sepharose CL6B beads (Cytiva). Elution profiles of antigen and BSA were monitored separately by fluorescence measurements of AF647 (ex/em: 650 nm / 665 nm) and AF488 (ex/em: 488 nm / 496 nm) on a multimodal plate reader (SpectraMax iD3, Molecular Devices, San Jose, CA, USA). Co-elution of antigen and albumin in the same fractions indicated albumin binding.

### Membrane insertion in B cells

Amphiphile insertion into cell membranes was assessed in vitro using the human-derived Ramos-RA1 B cell line (ATCC, CRL-1596). First, single cell suspensions were prepared and seeded at 5×10^6^ cells/ml (1×10^6^ cells per well) in a 96-well plate in cRPMI medium (RPMI-1640 supplemented with 10% fetal bovine serum (FBS) and 1% penicillin/streptomycin) containing 250, 500, 1000, 2000, or 3000 nM of AF647-labeled soluble protein or amph-protein for 1 hour at 37°C. After incubation, cells were washed once with PBS.

For quantification of membrane insertion by flow cytometry, cells were first stained with Live/Dead Aqua (Invitrogen) at 1:1000 dilution in 100 µl PBS for 15 minutes at 25°C. Cells were then washed once with FACS buffer (PBS + 1% BSA), then stained with AF555-VRC01 at 1.0 µg/10^6^ cells in 100 µl FACS buffer for 30 minutes at 4°C. Following staining, cells were washed twice with FACS buffer, then fixed with 2% paraformaldehyde and stored at 4°C until analysis. Samples were analyzed using a Symphony A3 flow cytometer.

For visualization of membrane insertion by confocal imaging, cells were stained with AF555-VRC01 at 1.0 µg/10^6^ cells in 100 µl FACS buffer for 30 minutes at 4°C, washed twice with FACS buffer, then fixed with 2% paraformaldehyde for 20-30 minutes at 25°C. 10 µl of fixed cell suspensions was loaded onto a 22 mm x 22 mm poly-L lysine-coated coverslip, then dried for 30–60 minutes in a humidified chamber. One drop of gold antifade mountant (Thermo Fisher Scientific) was added to a glass slide and the dried coverslip was placed onto the mountant. Excess mountant was removed, and the edges were sealed with clear nail polish to prevent drying. Slides were allowed to cure for 24 hours at room temperature before imaging using a Nikon A1Rsi HD Confocal microscope (UIC, University of Minnesota).

### In vitro B cell activation by calcium flux assay

To measure BCR-mediated B cell activation, calcium flux assays were performed in engineered germline VRC01 (glVRC01) B cells that express the human IgG1 broadly neutralizing monoclonal antibody VRC01, which is specific for the CD4 binding site of the HIV envelope glycoprotein gp120 (generously provided by Daniel Lingwood of the Ragon Institute) (*75*). Cells were first loaded with Fluo-8 calcium indicator dye: single-cell suspensions were seeded at 2×10^6^ cells/ml (0.2×10^6^ cells per well) in a black 96-well plate in equal parts HHBS buffer and Fluo-8 dye loading solution (Abcam, ab112129), then incubated for 1 hour at 37°C. Prior to reading each sample, the background fluorescence of each well (with glVRC01 cells and Fluo-8 dye) was recorded for ∼30 seconds. Next, different concentrations of soluble protein, amph-protein conjugates, or ‘painted’ cells (protein/amph-protein pre-incubated Ramos cells, prepared as described above) were added to the dye-loaded cells. Kinetic measurements of Fluo-8 fluorescence were then recorded in real-time at Ex/Em = 490/525 nm using a multimodal plate reader (SpectraMax iD3, Molecular Devices, San Jose, CA, USA) and plotted as an increase from the baseline fluorescence.

### Murine strains

All procedures were approved by the University of Minnesota Institutional Animal Care and Use Committee (IACUC). Procedures followed local, state, and federal regulations (protocol # 2407-42242A). Immunization studies were carried out using age-matched 6-8 week old female BALB/cJ mice (strain 000664) purchased from the Jackson Laboratory.

### Mouse immunizations and sample collection

Six-to-eight week old female BALB/cJ mice were immunized by subcutaneous tail base injection (100 μl total, split into 50 μl on each side of the tail) containing a 5 μg dose of protein (eOD or MD39) mixed with 25 μg cyclic di-GMP (cdGMP) adjuvant in PBS, as indicated. Animals were primed on day 0 and boosted on day 28 with an equal dose. For longitudinal immune monitoring, peripheral blood was collected bi-or tri-weekly via suborbital cheek bleed into Z-Gel serum separator microtubes (Cat# NC1029089, Thermo Fisher Scientific). Blood was allowed to clot at room temperature for 30 minutes, then serum was isolated by centrifuging at 10,000 × g for 5 minutes at 25°C. The resulting serum supernatant was collected and stored at −80 °C for subsequent ELISA analysis of antigen-specific IgG titers.

### IVIS trafficking in lymph nodes

To evaluate in vivo trafficking of AF647-labeled amph-proteins and proteins, BALB/c mice were immunized subcutaneously by tail base injection (as described above) and analyzed using an In Vivo Imaging System (IVIS Spectrum, UIC) for fluorescence detection. Mice were anesthetized and administered a 5 µg dose of AF647-amph-protein or AF647-protein mixed with 25 µg cdGMP adjuvant in PBS. Naïve mice served as controls. At 1 and 4 days post-immunization, mice were euthanized and inguinal lymph nodes (igLNs) were excised for fluorescence imaging. AF647 fluorescence (radiant efficiency) was quantified using In Vivo Imaging System (IVIS) fluorescence imaging (Living Image software, Perkin Elmer).

### Flow cytometry analysis of GC B cell activation

BALB/c mice were immunized by subcutaneous tail base injection (as described above) containing a 5 μg dose of protein (eOD or MD39) mixed with 25 μg cyclic di-GMP (cdGMP) adjuvant in PBS, as indicated. Mice were euthanized on day 12 and igLNs were isolated and mechanically digested by mashing in a 1.5ml biomasher tube (Kimble) for roughly 30 seconds in FACS buffer (PBS+1% BSA). The supernatant cell suspension was then passed through a 40µm cell strainer, centrifuged at 500<u>x</u>g for 5 minutes, washed twice and resuspended in FACS buffer. Cells were washed once with PBS and first stained with Live/Dead Aqua (Invitrogen) at 1:500 in 100 µl PBS for 15 minutes at 25°C, then treated with anti-mouse CD16/32 Fc block (TruStain FcX, BioLegend) at 1:100 in 50 µl FACS buffer for 10 minutes at 4°C. To identify Ag-specific germinal center (GC) B cells, half of the igLN cells from each mouse were stained with the following panel in 50µl FACS buffer for 30 minutes at 4°C: anti-mouse CD3ε BUV395 at 1:200 (clone 145-2C11, BioLegend), B220 AlexaFluor700 at 1:200 (RA3-6B2, BioLegend), CD38 BV421 at 1:200 (90, BioLegend), GL7 PE at 1:150 (GL7, BioLegend), eOD- or MD39-tetramer RB613 at 1:100, and eOD- or MD39-tetramer RB780 at 1:50. Fluorophore-labeled eOD- or MD39-tetramers were prepared by first reacting eOD or MD39 with maleimide-PEG2-biotin (Thermo Fisher Scientific) per the manufacturer’s instructions, then 5 molar equivalents of biotinylated-eOD or -MD39 were complexed with 1 molar equivalent of streptavidin-RB613 or streptavidin-RB780 (BD Biosciences), respectively, for 30 minutes at 25°C. To identify T follicular helper (Tfh) cells, half the igLN cells from each mouse were stained with the following antibodies in 50 µl FACS buffer for 30 minutes at 4°C: anti-mouse B220 BV510 at 1:200 (clone RA3-6B2, BioLegend), CD4 BV711 at 1:200 (GK1.5, BioLegend), CD44 PE-Cy7 at 1:200 (IM7, BioLegend), Inducible T Cell Costimulator (ICOS) PE at 1:100 (7E.17G9, BioLegend), programmed cell death protein 1 (PD-1) BV650 at 1:50 (J43, BD Biosciences), and CXCR5-biotin at 1:50 (2G8, BD Biosciences) followed by streptavidin-BV421 at 1:100 (BioLegend).

### ELISA analyses of murine antibody titers

Antigen-specific IgG titers were measured in mouse serum samples by enzyme-linked immunosorbent assay (ELISA). To capture eOD-specific antibodies, Costar Polystyrene High Binding 96-well plates (Corning) were coated with 1 µg/mL eOD antigen in PBS and incubated overnight at 4°C, then blocked with blocking buffer (PBS + 2% BSA) for 2 hours at 25°C. To capture MD39-specific antibodies, Costar Polystyrene High Binding 96-well plates (Corning) were coated with 10 µg/mL streptavidin in PBS overnight at 4°C, blocked with PBS containing 2% BSA for 2 hours at 25°C, then coated with biotinylated-MD39 at 2 µg/mL in blocking buffer overnight at 4°C. Separately, mouse serum samples were diluted in blocking buffer, starting at 1:100, followed by 4-fold serial dilutions. hVRC01 (5 µg/mL) was included as a positive control. Serially diluted samples were incubated in coated plates for 2 hours at 25°C, followed by incubation with 1:5000 goat anti-mouse IgG-HRP (Bio-Rad) for mouse samples or 1:5000 goat anti-human IgG-HRP (Invitrogen) for positive control VRC01 in block buffer for 1 hour at 25°C. Plates were developed using TMB substrate (3,3′,5,5′-tetramethylbenzidine) for 1–20 minutes and the reaction was stopped with 2N sulfuric acid. Absorbance was measured at 450 nm with 540 nm background subtraction (A450–A540) on a multimodal plate reader (SpectraMax iD3, Molecular Devices, San Jose, CA, USA). For all comparisons, samples were developed for the same duration to ensure consistency. Cutoff titers were determined as the inverse dilution giving an HRP absorbance of 0.1 based on background.

### ELISPOT analysis of murine plasma cells and memory B cells

Total and antigen specific IgG plasma cell responses in bone marrow (BM) were measured at 22 weeks post prime using 96-well multiscreen HTS filter plates (Millipore) and mouse IgG/IgA ELISPOT Flex kits (MABTECH). Filter plates were coated overnight at 4°C with 100 µl/well of 15 µg/mL anti-IgG (for total IgG ASC detection) and 1 µg/mL eOD/MD39 (for Ag-specific ASC detection) . Each plate was washed 5 times with 200 µL/well of PBS and then blocked with 200 µL/well of cRPMI. Cells were plated at 500,000 and 250,000 cells per well and incubated at 37°C and 5% CO 2 for 16 hours. Spot detection was carried out per manufacturer instructions, and plates were read on a CTL ImmunoSpot Analyzer.

### Statistical analysis

Statistics were analyzed using GraphPad Prism software. For comparison of two groups, a two-tailed unpaired t test was performed. For comparisons of more than two groups, a one- or two-way analysis of variance (ANOVA) was performed with α = 0.05, followed by Tukey’s or Sidak’s post hoc test for multiple comparisons as appropriate. Statistical significance in the MFI plots of the cell-painting experiment was determined using simple linear regression to evaluate whether the dependence of AF647 or VRC01 mean fluorescence intensity on antigen concentration resulted in a significant non-zero slope. All graphs represent means ± SEM unless otherwise noted. Statistical significance is denoted as: *p < 0.05, **p < 0.01, ***p < 0.001, and ****p < 0.0001.

## Acknowledgements

We thank William Schief and Darrell Irvine who generously provided eOD-gp120 and MD39 proteins and plasmids, and Mariane Melo for her guidance with protein production. We gratefully acknowledge the staff and resources with the University of Minnesota Research Animal Resources (RAR) for providing animal support and expertise, as well as the University Imaging Centers (UIC), the UMN Characterization Facility, UMN Flow Core (UFCR), and Minnesota Nano Center. The UMN Characterization Facility receives partial support from the NSF through the MRSEC (Award Number DMR-2011401) and the NNCI (Award Number ECCS-2025124) programs. The Nano Center is supported by the National Science Foundation through the National Nanotechnology Coordinated Infrastructure (NNCI) under Award Number ECCS-2025124. Figure schematics were created with Biorender.

## Author contributions

D.Y. and B.L.H. conceptualized the study. D.Y., E.L.T., and K.J. contributed to the methodology. D.Y., K.J., M.L.S., B.H., J.M.L., and N.S. carried out the experimental investigation. D.Y. and E.L.T. produced eOD and MD39. D.Y. synthesized and characterized amph-PEG2K-eOD, amph-PEG5K-eOD, and amph-MD39. D.Y. performed mouse immunization studies, ELISPOT, flow cytometry, confocal microscopy, cell painting experiments, and albumin binding assays. D.Y. and K.J. performed the calcium-flux assays. D.Y. and M.L.S. performed serum sample collection and ELISAs. D.Y. performed antigenicity ELISAs. D.Y., B.H., and J.M.L. performed IVIS. D.Y., E.L.T., M.L.S., B.H., J.M.L., and N.S. performed tissue harvests and processing for IVIS, flow cytometry, and ELISPOT. D.Y. performed data analysis. B.L.H. contributed to project administration. D.Y. and B.L.H. were involved with reviewing and editing the draft.

## Data availability

All data associated with this study are in the paper or the Supplementary Materials.

