## Supplementary Materials for "‘Cell painting’ with amphiphile-protein conjugate vaccines drives B cell activation and germinal center priming to enhance humoral immunity"

### Supplemental Figures

**A** **eOD-PADRE:** MW 21.787 kDa
ETG**C**HHHHHHGGDTITLPCRPAPPPHCSSNITGLILTRQGGYSNDNTVIFRPSGGDWRDIARCQIAGTVVSTQLFLNG
SLAEEEVVIRSEDWRDNAKSICVQLNTSVEINCTGAGHCNISRAKWNNTLKQIASKLREQYGNKTIIFKPSSGGDPEFV
NHS FNCGGEFFYCDSTQLFNSTWFNSTGS**AKFVAAWTLKAAA**

**B** **MD39 SOSIP trimer:** MW 217.018 kDa
Monomer sequence: (monomer MW 72.339 kDa)
AENLWVTVYYGVPVWKDAETTLFCASDAKAYETEKHNVWATHACVPTDPNPQEIHLENVTEEFNMWKNNMVEQMH
EDIISLWDQSLKPCVKLTPLCVTLQCTNVTNNITDDMRGELKNCSFNMTTELRDKKQKVYSLFYRLDVVQINENQGNR
SNNSNKEYRLINCNTSAITQACPKVSFEPIPIHYCAPAGFAILKCKDKKFNGTGPCPSVSTVQCTHGIKPVVSTQLLLNG
SLAEEEVIIRSENITNNAKNILVQLNTPVQINCTRPNNNTVKSIRIGPGQAFYYTGDIIGDIRQAHCNVSKATWNETLGKV
VKQLRKHFGNNTIIRFAQSSGGDLEVTTHSFNCGGEFFYCNTSGLFNSTWISNTSVQGSNSTGSNDSITLPCRIKQIINM
WQRIGQAMYAPPIQGVIRCVSNITGLILTRDGGSTNSTTETFRPGGGDMRDNWRSELYKYKVVKIEPLGVAPTRCKRR
VVGRRRRRRAVGIGAVSLGFLGAAGSTMGAASMTLTVQARNLLSGIVQQQSNLLRAPEPQQHLLKDTHWGIKQLQAR
VLAVEHYLRDQQLLGIWGCSGKLICCTNVPWNSSWSNRNLSEIWDNMTWLQWDKEISNYTQIIYGLLEESQNQQEKN
EQDLLALDGTKHHHHHH**C**

**C**

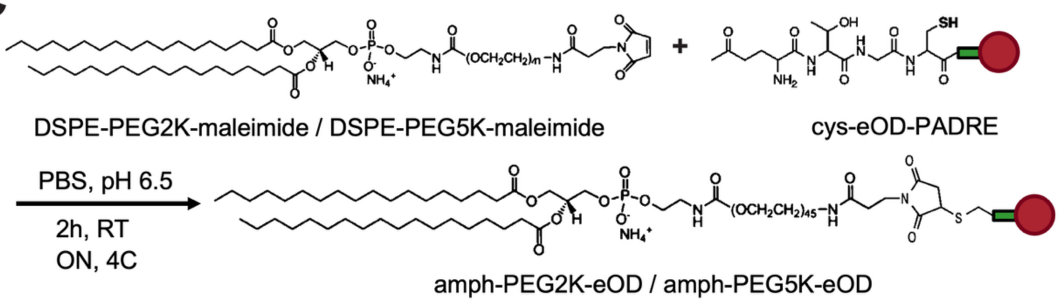


**D**

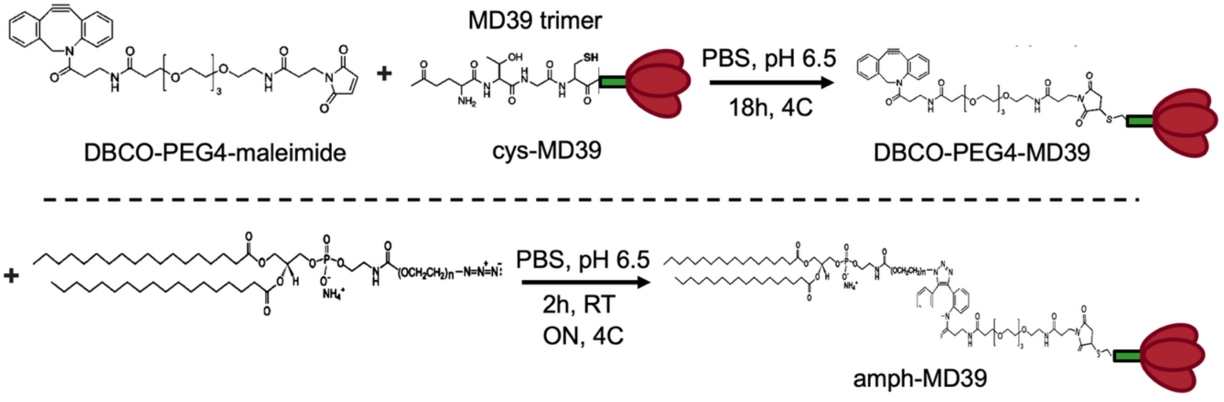


**Figure S1. Synthesis of amphiphile-protein conjugates. A)** eOD-PADRE protein sequence with pan HLA DR-binding epitope (PADRE) peptide shown in red. **B)** MD39 protein sequence, showing monomer. **C)** Reaction scheme for preparation of amph-eOD conjugates and **D)** amph-MD39 conjugates**.**

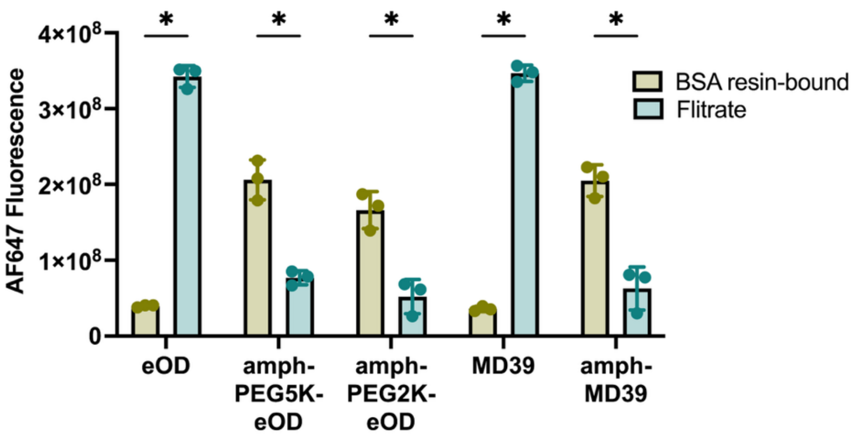


**Figure S2. Amphiphile conjugation of eOD and MD39 protein antigens enables albumin binding.** AF647 fluorescence signal of albumin-conjugated agarose resin compared to the filtrate after incubation of AF647-labeled conjugates with BSA-resin for 1 hr at 37 °C, followed by purification by affinity chromatography.

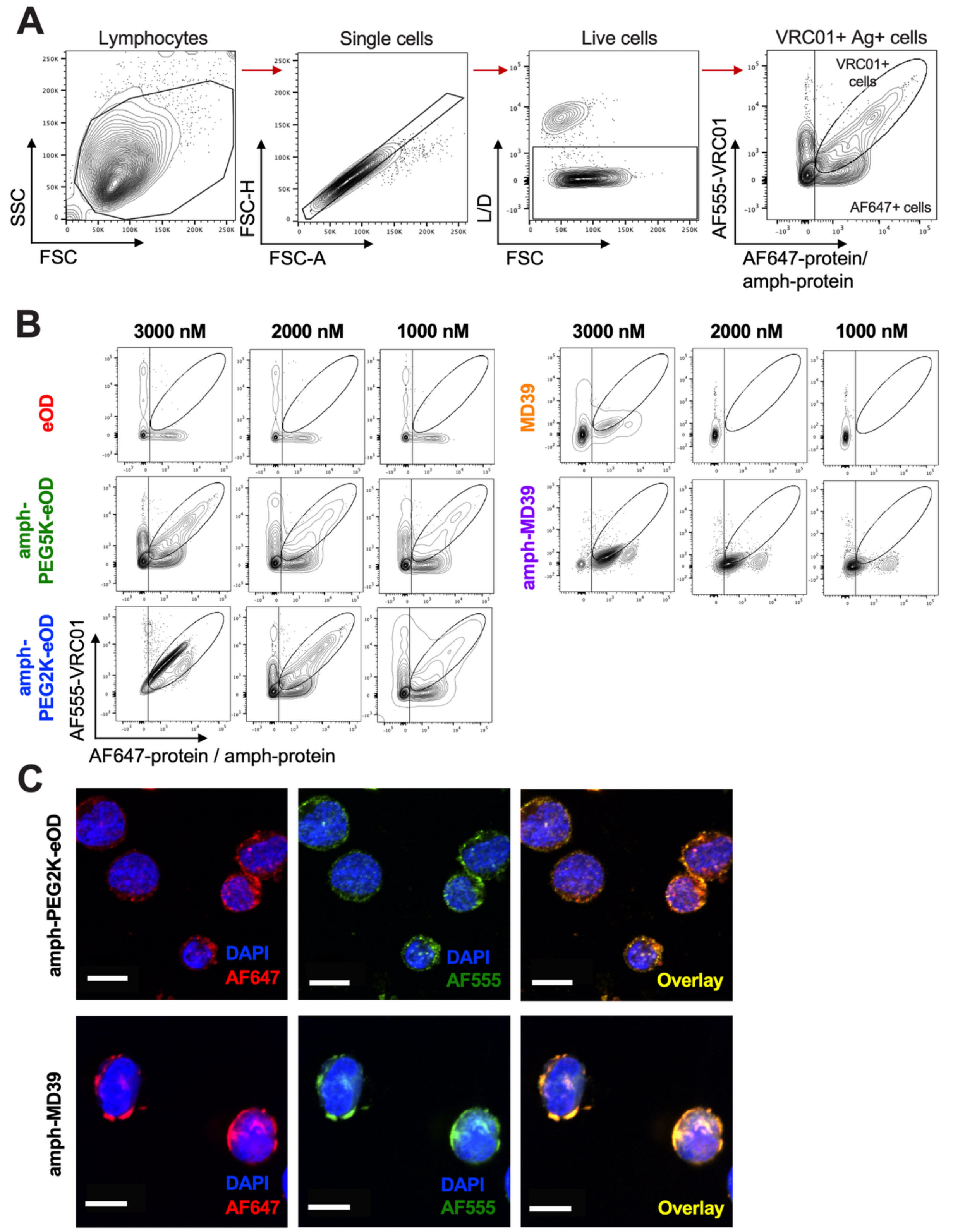


**Figure S3. Cell membrane insertion assays.** Ramos B cells were incubated with AF647-labeled protein or amphiphile-protein conjugates (1000, 2000, or 3000 nM) for 1 hour at 37°C. Cells were then washed and stained with AF555-labeled VRC01 antibody. Membrane insertion (cell painting)

was measured by flow cytometry. **A)** The gating strategy is shown for identification of VRC01$^+$ and AF647$^+$ cells; FSC, forward scatter; SSC, side scatter; A, area; H, height; L/D, live/dead dye. **B)** Representative flow cytometry plots of protein / amph-protein and VRC01 binding to the cells at varying concentrations of protein (eOD or MD39). FBS, fetal bovine serum. **C)** Representative images obtained by confocal microscopy showing the localization of AF647-labeled immunogens (red) in Ramos B cells stained with DAPI (blue) and AF555-VRC01 (green) on the cell membrane. Scale bars equals 10 μm (60X).

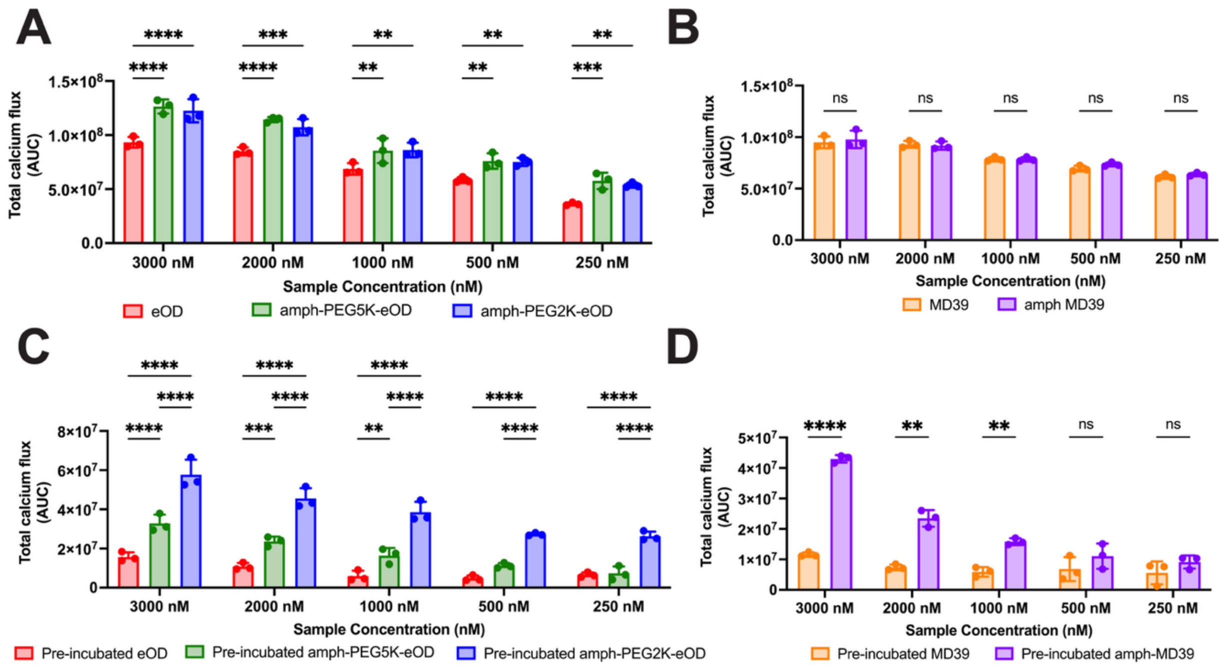


**Figure S4. Amphiphile-protein conjugates enhance B cell activation**. glVRC01 B cells were loaded with a calcium indicator dye (Fluo-8) to monitor BCR signaling induced by protein or amph-protein conjugates. Activation was measured in real-time as Fluo-8 signal upon addition of immunogens on a fluorescent plate reader. In a separate independent experiment, Ramos B cells were first pre-incubated with immunogens for 1 hr at 37°C to allow for membrane insertion, then washed Ramos cells were added to dye-loaded glVRC01 cells. **A-B)** Total calcium flux (quantified as integrated area under the curve, AUC) from glVRC01 cell incubation with **A)** eOD and **B)** MD39 *constructs* at all tested concentrations. **C-D)** Total calcium flux AUC from glVRC01 cell incubation with **C)** eOD- and **D)** MD39-*painted Ramos cells* at all tested concentrations. Statistical significance determined by ordinary one-way ANOVA followed by Tukey's post hoc test (for eOD groups) or unpaired t-test (for MD39 groups). All data shown are presented as mean ± SEM (n=3).

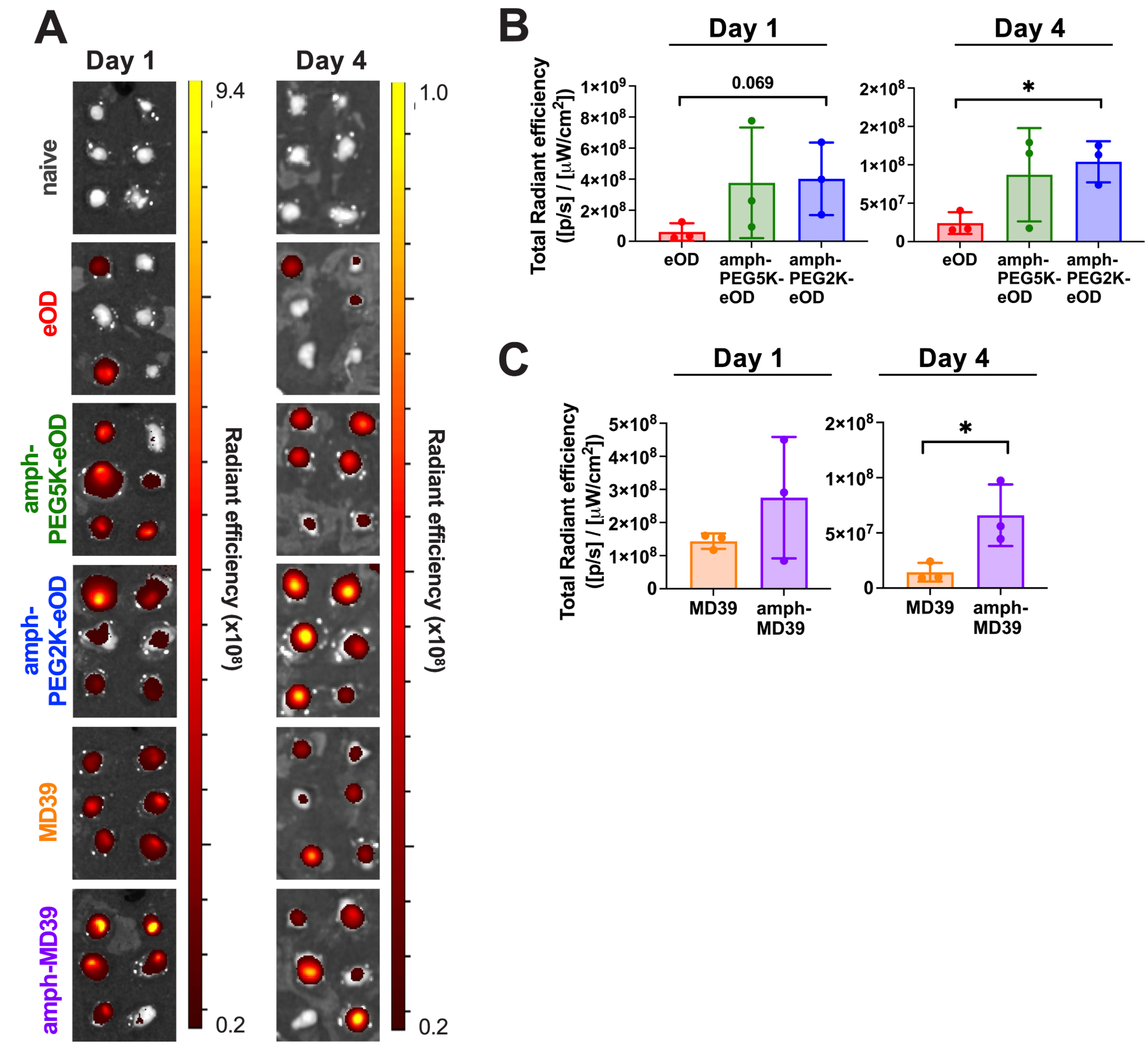


**Figure S5. Vaccine uptake and retention in the inguinal lymph nodes.** Groups of Balb/c mice (n=3 animals per group) were immunized subcutaneously (s.c.) at the tail base with AF647-labeled protein or amph-protein conjugates mixed with cdGMP adjuvant. Inguinal lymph nodes (igLNs) were isolated and imaged by IVIS at one and four days post-immunization to measure vaccine uptake. **A)** Representative IVIS images of vaccine AF647 signal in all respective igLNs over time following s.c. administration. **B-C)** Quantified IVIS signal from (A) in igLNs at day 1 and 4, shown as total radiant efficiency. p, photon. Statistical significance determined at each time point by ordinary one-way ANOVA followed by Tukey's post hoc test (for eOD groups) or unpaired t-test (for MD39 groups). All data presented as mean ± SEM.

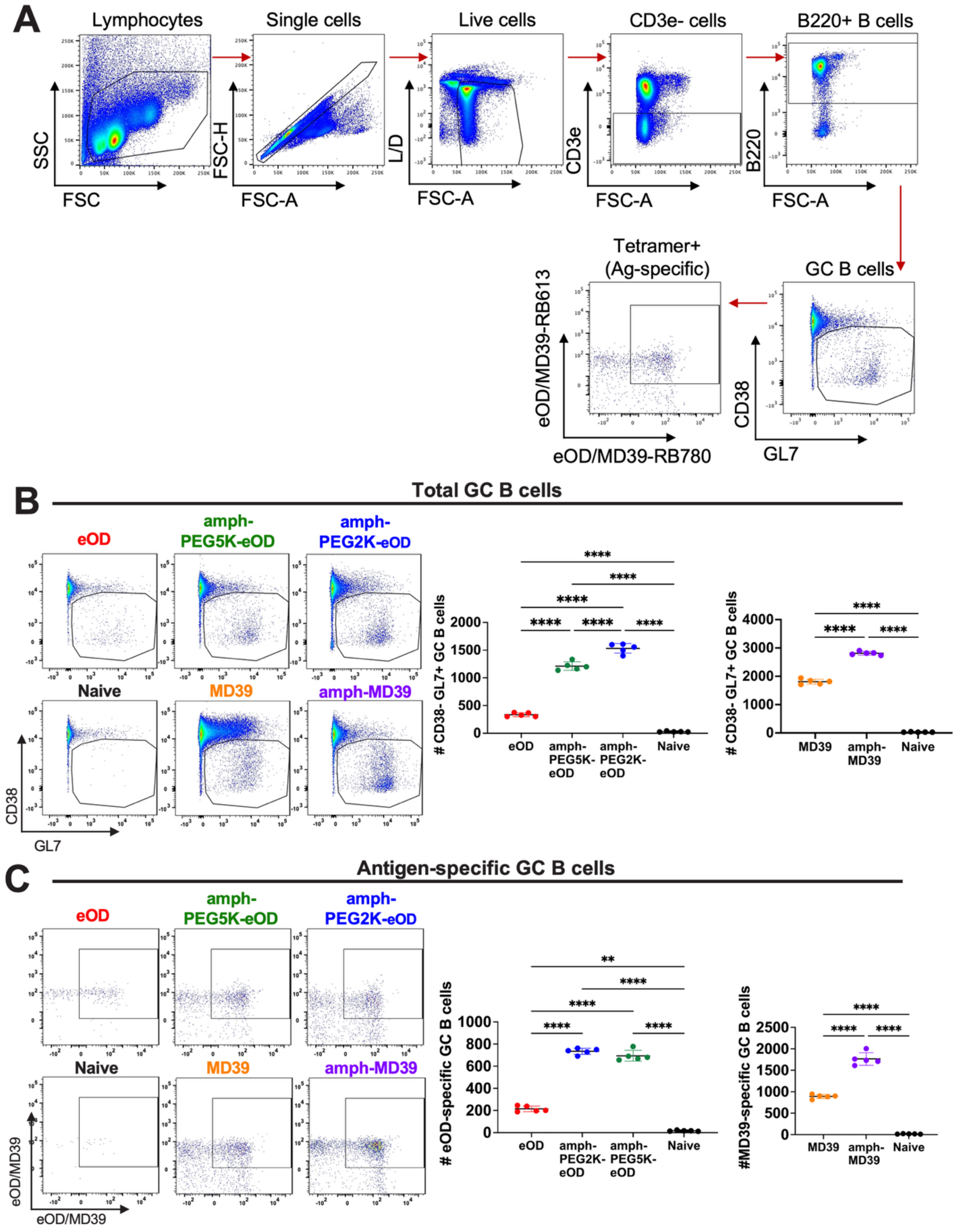


**Figure S6. GC B cell responses in mouse igLNs following s.c. immunization with amph-protein.** Groups of BALB/c mice (n = 5 animals per group) were immunized with 5 μg protein or amph-protein conjugates mixed with 5 μg cdGMP adjuvant, and GC responses were analyzed by flow cytometry on day 12. **A)** The gating strategy for identification of GC B cells is shown. **B)** Representative flow cytometry plots and absolute number of cells are presented, showing total $CD38^{-}GL7^{+}$ GC B cells for all igLN samples, including controls. **C)** Representative flow cytometry plots and absolute number of cells are presented, showing eOD- or MD39-tetramer$^{+}$ GC B cells for all igLN samples, including controls. Statistical significance was determined by ordinary one-way ANOVA followed by Tukey's post hoc test. All data presented as mean ± SEM. *$p$<0.05, **$p$<0.01, ***$p$<0.001, and ****$p$<0.0001.

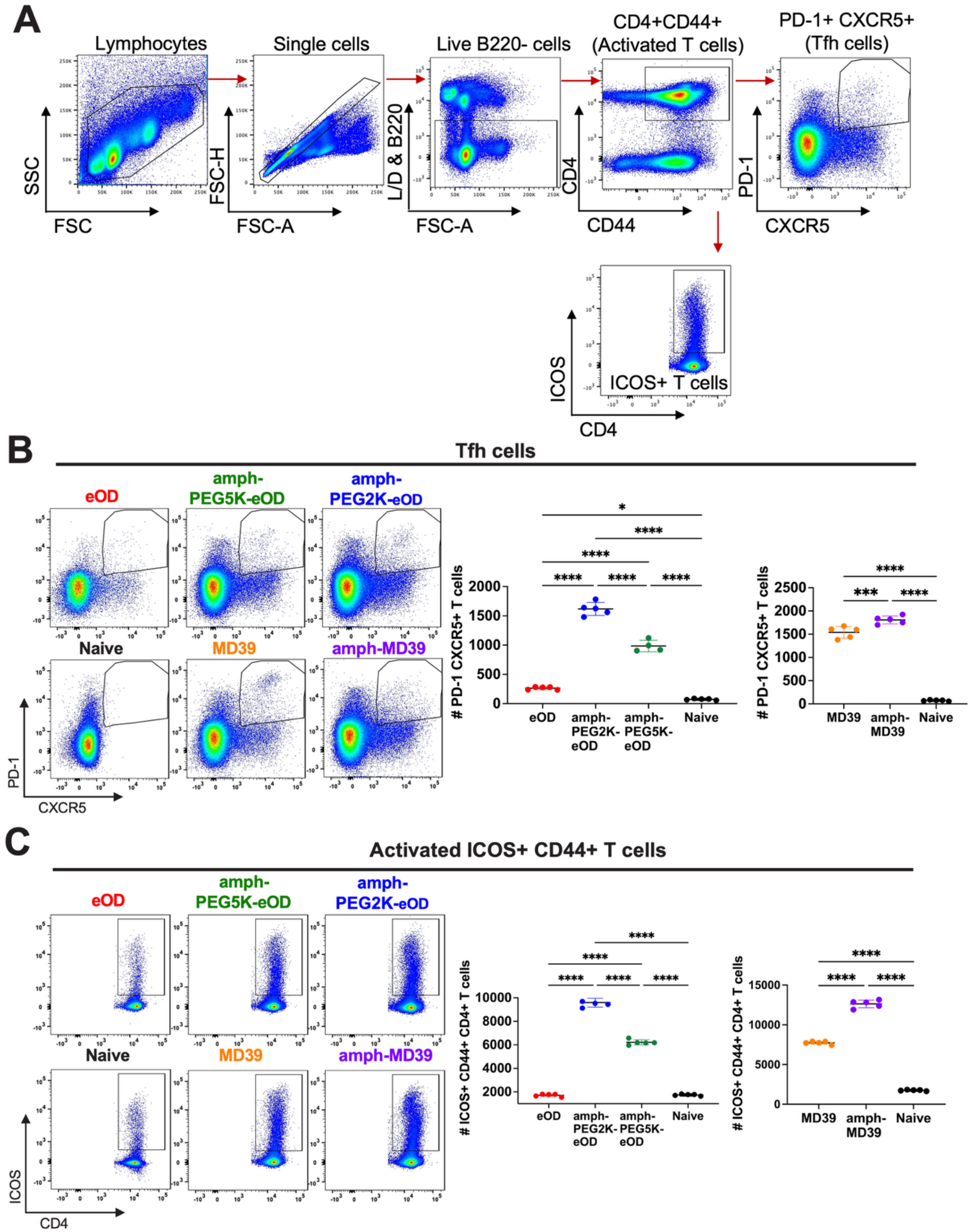


**Figure S7. Tfh cell responses in mouse igLNs following s.c. immunization with amph-protein.** Groups of BALB/c mice (n = 5 animals per group) were immunized with 5 μg protein or amph-protein mixed with 5 μg cdGMP adjuvant, and Tfh responses were analyzed by flow cytometry on day 12. **A)** The gating strategy for identification of Tfh cells is shown. **B)** Representative flow cytometry plots and absolute number of cells are presented, showing total PD-1$^{+}$CXCR5$^{+}$ Tfh cells for all igLN samples, including controls. **C)** Representative flow cytometry plots and absolute number of cells are presented, showing ICOS$^{+}$CD4$^{+}$CD44$^{+}$ T cells for all igLN samples, including controls. Statistical significance was determined by ordinary one-way ANOVA followed by Tukey's post hoc test. All data presented as mean ± SEM. *$p$<0.05, **$p$<0.01, ***$p$<0.001, and ****$p$<0.0001.
